# Youthful and age-related matreotypes predict drugs promoting longevity

**DOI:** 10.1101/2021.01.26.428242

**Authors:** Cyril Statzer, Elisabeth Jongsma, Sean X. Liu, Alexander Dakhovnik, Franziska Wandrey, Pavlo Mozharovskyi, Fred Zülli, Collin Y. Ewald

## Abstract

The identification and validation of drugs that promote health during aging (‘geroprotectors’) is key to the retardation or prevention of chronic age-related diseases. Here we found that most of the established pro-longevity compounds shown to extend lifespan in model organisms also alter extracellular matrix gene expression (*i.e.,* matrisome) in human cell lines. To harness this novel observation, we used age-stratified human transcriptomes to define the age-related matreotype, which represents the matrisome gene expression pattern associated with age. Using a ‘youthful’ matreotype, we screened *in silico* for geroprotective drug candidates. To validate drug candidates, we developed a novel tool using prolonged collagen expression as a non-invasive and *in-vivo* surrogate marker for *C. elegans* longevity. With this reporter, we were able to eliminate false positive drug candidates and determine the appropriate dose for extending the lifespan of *C. elegans*. We improved drug uptake for one of our predicted compounds, genistein, and reconciled previous contradictory reports of its effects on longevity. We identified and validated new compounds, tretinoin, chondroitin sulfate, and hyaluronic acid, for their ability to restore age-related decline of collagen homeostasis and increase lifespan. Thus, our innovative drug screening approach - employing extracellular matrix homeostasis - facilitates the discovery of pharmacological interventions promoting healthy aging.

**Highlights:**

- Many geroprotective drugs alter extracellular matrix gene expression
- Defined young and old human matreotype signatures can identify novel potential geroprotective compounds
- Prolonged collagen homeostasis as a surrogate marker for longevity

## Introduction

The demographic shift in the human population reflects an ageing society - over 20% of Europeans are predicted to be 65 or over by the year 2025 (Riera and Dillin, 2015). Aging is the major risk factor for developing chronic diseases, such as cancer, Alzheimer’s disease, and cardiovascular complications (Niccoli and Partridge, 2012). Unfortunately, humans spend on average one-fifth of their lifetime in poor health suffering from one or multiple age-related chronic diseases (Partridge et al., 2018). However, the onset of age-related pathologies is not fixed, and the rate of aging was shown to be malleable. The goal of biomedical research on aging or geroscience is to identify interventions that compress late-life morbidity to increase the period spent healthy and free from disease (Barzilai et al., 2018; Campisi et al., 2019; Kennedy et al., 2014; Olshansky, 2018; Partridge et al., 2018; Riera and Dillin, 2015).

A few geroprotective drugs exist that postpone age-related diseases (Riera and Dillin, 2015). For instance, the anti-diabetes drug metformin reduces age-related chronic diseases and mortality from all causes (Bannister et al., 2014; Campbell et al., 2017). Ongoing clinical trials on geroprotective drugs or compounds include the anti-diabetic drugs metformin (NCT02432287, NCT03451006) (Barzilai et al., 2016) and acarbose (NCT02953093); mTOR-inhibiting and immunosuppressant drug rapamycin (sirolimus; NCT02874924) (Mannick et al., 2014, 2018); natural compounds resveratrol (NCT01842399) and Urolithin A (NCT04160312) (Andreux et al., 2019); and nicotinamide adenine dinucleotide precursors NR (NCT02950441) and NMN (NCT04685096) (Tsubota, 2016). Although clinical trials targeting aging are challenging due to a lack of clear biomarkers of the aging process (Espeland et al., 2017), one primary outcome measure used in the aforementioned clinical trials for metformin and acarbose is the restoration from an ‘old’ to a ‘youthful’ gene expression signature (NCT02432287, NCT02953093). Therefore, we reasoned that cross-comparing youthful expression signatures against expression profiles elicited by small molecules could identify novel geroprotective compounds. Conceptually similar *in-silico* approaches have been conducted in the past (Aliper et al., 2016; Calvert et al., 2016; Dönertaş et al., 2018, 2019; Fuentealba et al., 2019; Janssens et al., 2019; Komljenovic et al., 2019; Liu et al., 2016). However, these former approaches comparing age-related expression profiles with drug-treated cells revealed similar drug targets (Dönertaş et al., 2019), such as the HSP90/HSF-1 axis (Fuentealba et al., 2019; Janssens et al., 2019). Our strategy was to use a more refined starting list of a ‘youthful’ gene expression signature with experimentally implicated genes associated with healthy aging.

A key signature of aging is the continuous decline of collagen and cell adhesion gene expression (Ewald, 2019; Magalhães et al., 2009; Zhavoronkov et al., 2014) accompanied with an increase in matrix metalloproteinase expression (Ewald, 2019; Fisher et al., 2009). Gene expression ontologies of extracellular matrix (ECM) genes have been associated with healthy aging in humans (Zeng et al., 2020). The ECM not only embeds cells and tissues but also provides instructive cues that change cellular function and identity. For instance, placing old cells into a ‘young’ ECM rejuvenates senescent cells or stem cells and even reprograms tumor cells (reviewed in (Ewald, 2019)). Moreover, collagen homeostasis is required and sufficient for longevity in *C. elegans* (Ewald et al., 2015). Heparan/chondroitin biosynthesis and TGFβ pathway are frequently enriched in *C. elegans* longevity drug screens (Liu et al., 2016). Collectively taken, these functionally implicated genes are all members of the matrisome.

The human matrisome encompasses 1027 protein-encoding genes that either form the ECM, such as collagens, glycoproteins, and proteoglycans; associate with ECM (*e.g.,* TGFβ, Wnts, cytokines); or remodel the ECM (*e.g.,* matrix metalloproteinases (MMPs)) (Naba et al., 2016). The matrisome represents about 4% of the human genome and is functionally implicated in about 8% of the total 7037 unique human phenotypes (Statzer and Ewald, 2020). Age-related phenotypes rank among the top matrisome-phenotypic categories across species (Ewald, 2019; Taha and Naba, 2019). Proteomics approaches have revealed unique ECM compositions associated with health and disease status (Socovich and Naba, 2019) — ECM compositions can even be used to identify distinct cancer-cell types (Ewald, 2019). Therefore, organismal phenotypes, physiological states, and cellular identity are characterized by distinct sets of expressed ECM proteins. Since these unique ECM compositions are an expression profile on a temporary, sometimes local basis and do not involve the entire matrisome, we coined the term matreotype (Ewald, 2019). The matreotype is the acute state of an ECM composition associated with- or causal for- a given physiological condition or phenotype (Ewald, 2019).

Given the functional implication of ECM in healthy aging, we hypothesized that a youthful matreotype might predict drugs promoting healthy aging. Here we define a youthful human matreotype using data from the Genotype-Tissue Expression (GTEx) project (Consortium, 2013). We query this young matreotype signature with the drug resource Connectivity Map (CMap) (Lamb et al., 2006) data to identify longevity-promoting compounds. We then developed a novel *in-vivo* tool as a surrogate marker for longevity to find appropriate drug doses to be used for *C. elegans’* lifespan assays. Our results implicate previously known longevity drugs as well as novel drugs, providing a proof-of-concept for our approach.

## Results

### Geroprotective compounds associated with altering ECM

We first applied literature and database mining to search for compounds that have been shown to increase lifespan and are known to alter ECM in any organism. We acquired lifespan data from databases DrugAge and GeroProtectors (Figure 1A) (Barardo et al., 2017a; Moskalev et al., 2015). We filtered for reported mean lifespan extensions that were above 5% for compounds compared to control (Figure 1A; Supplementary Table 1)—then queried all PubMed abstracts for the given agent and our ECM key terms, including collagen, MMP, proteoglycan, integrin, TGFβ (Figure 1A; Supplementary Table 1). After manual curation, we identified approximately 3% (16 out of 567) of the examined known longevity-promoting compounds that slow aging and also had been reported to affect proteins outside of cells, such as collagens and other matrisome proteins (Figure 1B; Supplementary Table 1).

**Figure 1.**
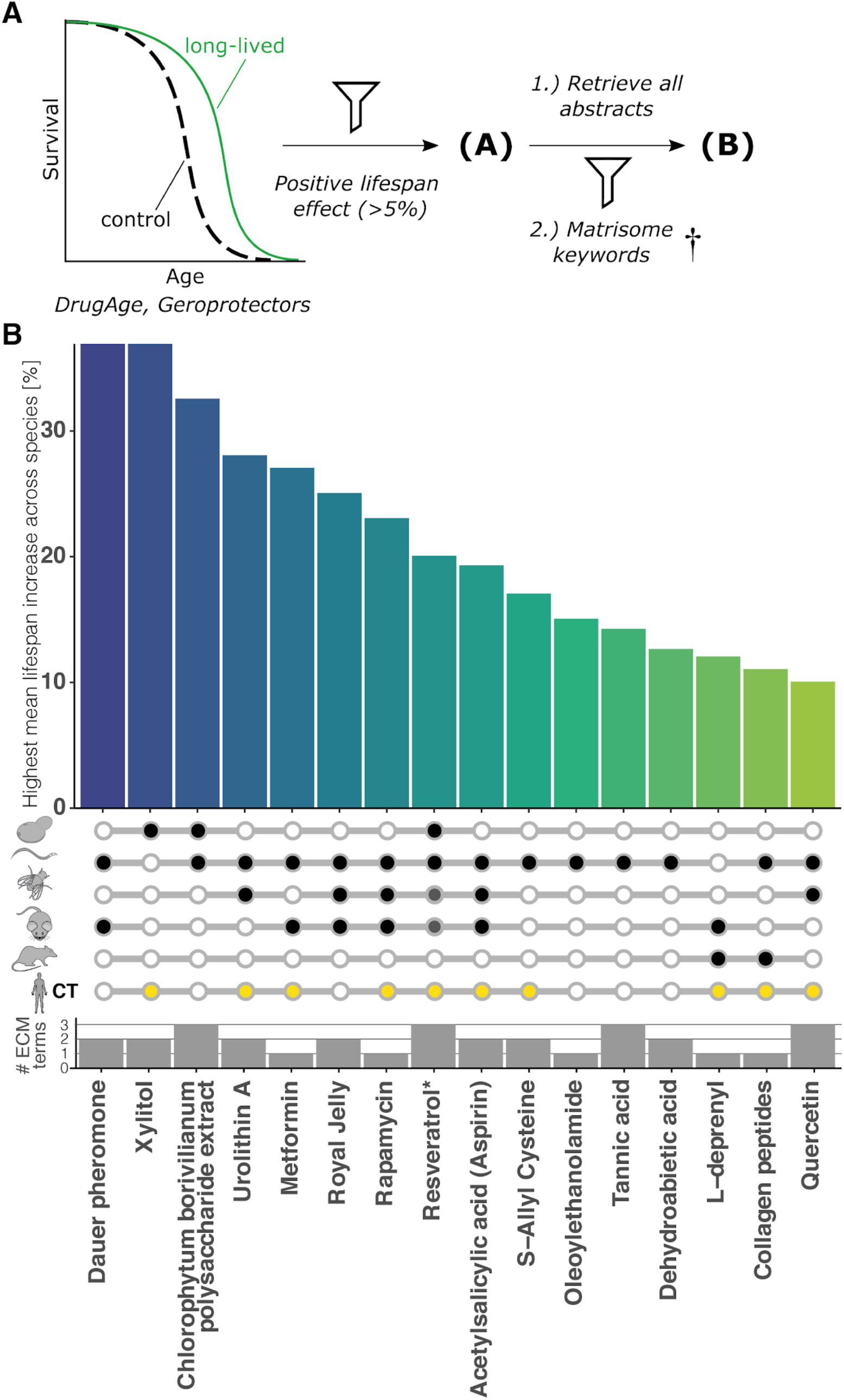
Geroprotective compounds shown to affect the ECM. (A) Schematic depiction of the literature and database mining approach used. (B) Compounds which contained ECM keywords in their abstract were ranked by their highest mean lifespan increase in any species. Open circle = not assessed, black closed circle = lifespan increased, yellow circle = compound assessed in clinical trials (CT). The number of ECM keywords quantified in the abstract is displayed below. *Resveratrol only on a high-fat diet increased lifespan in fly and mouse. For details and references, see Supplementary Table 1.

### Longevity compounds affect matrisome gene expression

Next, we investigated whether compound treatments, in general, would alter matrisome expression. Connectivity Map (CMap) is a library of 1.5 million gene expression profiles comparing 1309 different compound treatments on human cell cultures (Lamb et al., 2006). We queried the CMap library for compound treatments that either increase or decrease the expression levels of matrisome genes. Using a z-score threshold of ± 1.5, we identified 167 compounds that strongly regulate the 594 out of the 1027 matrisome genes compared to the background (13752 total quantified genes; Figure 2A, Supplementary Figure 1, Supplementary Table 2). Out of the 12 most up and 12 most down-regulated matrisome expression profiles upon a compound treatment, we identified ten agents linked to longevity or impairment of age-related pathologies (Figure 2B, 2C, Supplementary Table 2). Strikingly, out of the 47 known longevity-promoting compounds that were assessed in the CMap library, we found 41 (87%) significantly altered matrisome gene expression (Supplementary Figures 2-4, Supplementary Table 2). After manual curation, we identified 20 additional compounds reported to increase lifespan. From the total 67 reported lifespan-increasing compounds, 19 had minor effects on overall matrisome gene expression, whereas 26 compounds increased, and 22 decreased, matrisome gene expression (Figure 2A, Supplementary Table 2). These results suggest that compounds implicated in healthy aging show enriched differential matrisome gene expression.

**Figure 2.**
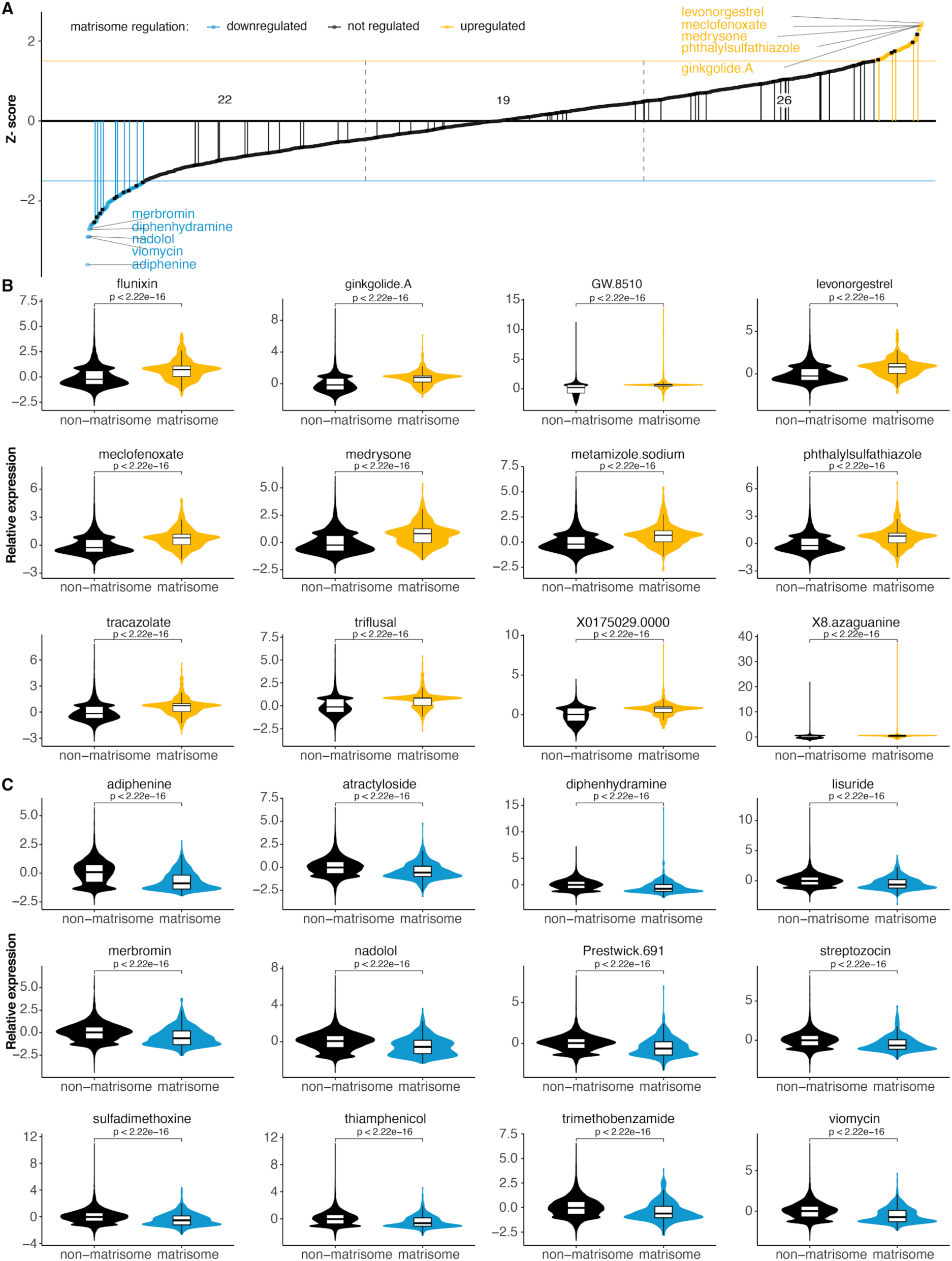
Drugs affecting matrisome gene expression. (A) Z-score of altering matrisome gene expression across all 1309 CMap drugs. Vertical lines indicate compounds shown to increase lifespan in any organism. (B-C) Top 12 compounds that increase (B) or decrease (C) the overall matrisome gene expression. For details, see Supplementary Table 2.

### Defining young and old human matreotypes

For a more targeted approach, we reasoned that using a young age-associated matrisome gene expression signature (*i.e.,* youthful matreotype) to query CMap data should reveal more compound treatments that might promote longevity. To define a young and old human matreotype, we built upon the previous analysis of the GTEx dataset (Consortium, 2013) comparing a young versus an old gene expression pattern of more than 50 tissues and 8000 transcriptomes by Janssens and colleagues (Janssens et al., 2019). From these 50 tissues, we only identified 15 tissues that had on average 138 age-related transcripts per tissue, which we then filtered for matrisome genes (Supplementary Figures 5, 6, Supplementary Table 3). In our analysis, we used two different approaches to quantify age-related transcript changes: the difference in expression (’absolute’) and the fraction of change (’relative’). The absolute expression change preferentially captures genes exhibiting high baseline expression levels since any change in their abundance translates to a large absolute change. By contrast, the relative expression difference quantifies the change to the previously measured value and favors lower expressed genes.

Among these age-related transcripts, matrisome genes were well represented with collagen genes and matrix proteases (MMPs, ADAMs) being decreased and increased, respectively, with age (Figure 3A-D, Supplementary Table 3). These findings were consistent with previous reports (Ewald, 2019; Fisher et al., 2009; Magalhães et al., 2009). However, we noted a number of novel observations. Firstly, each tissue has a unique age-related tissue-specific matreotype gene expression signature (Supplementary Figures 5, 6, Supplementary Table 3). Secondly, both the transcript coverage and the age-association of the matrisome varies within the 15 assessed tissues. Therefore, we decided to narrow the focus of our study to five tissues: skin, thyroid, pituitary, aorta, and coronary artery tissue (Supplementary Figure 7, Supplementary Table 3). In the remaining tissues, the number of age-associated transcripts was too low to quantify the contribution of the matrisome conclusively. Thirdly, certain matrisome genes, such as GDF15, experienced both increasing and decreasing expression levels during aging, depending on the tissue (Figure 3B). With these observations in mind, our aim was to construct a multi-tissue compendium of matrisome members, which were most affected by aging. To build upon our five tissues, we combined the findings of eight studies that also included transcriptomes across ages from different tissues in order to validate and identify additional multi-tissue and age-related matrisome genes (Supplementary Table 3). By considering both absolute and relative aging-gene expression changes, we determined the age-related matreotype of brain, fat, skin, and other tissues to generate a new common signature across tissues (Supplementary Figure 8, Supplementary Table 3). By inverting the aged expression pattern, we defined the age-reversed or ‘youthful’ matreotype (Supplementary Table 4). Using this approach, we defined here, for the first time, a multi-tissue compendium of approximately 100 genes across 15 human tissues, which we define as the young and aged matreotype.

**Figure 3.**
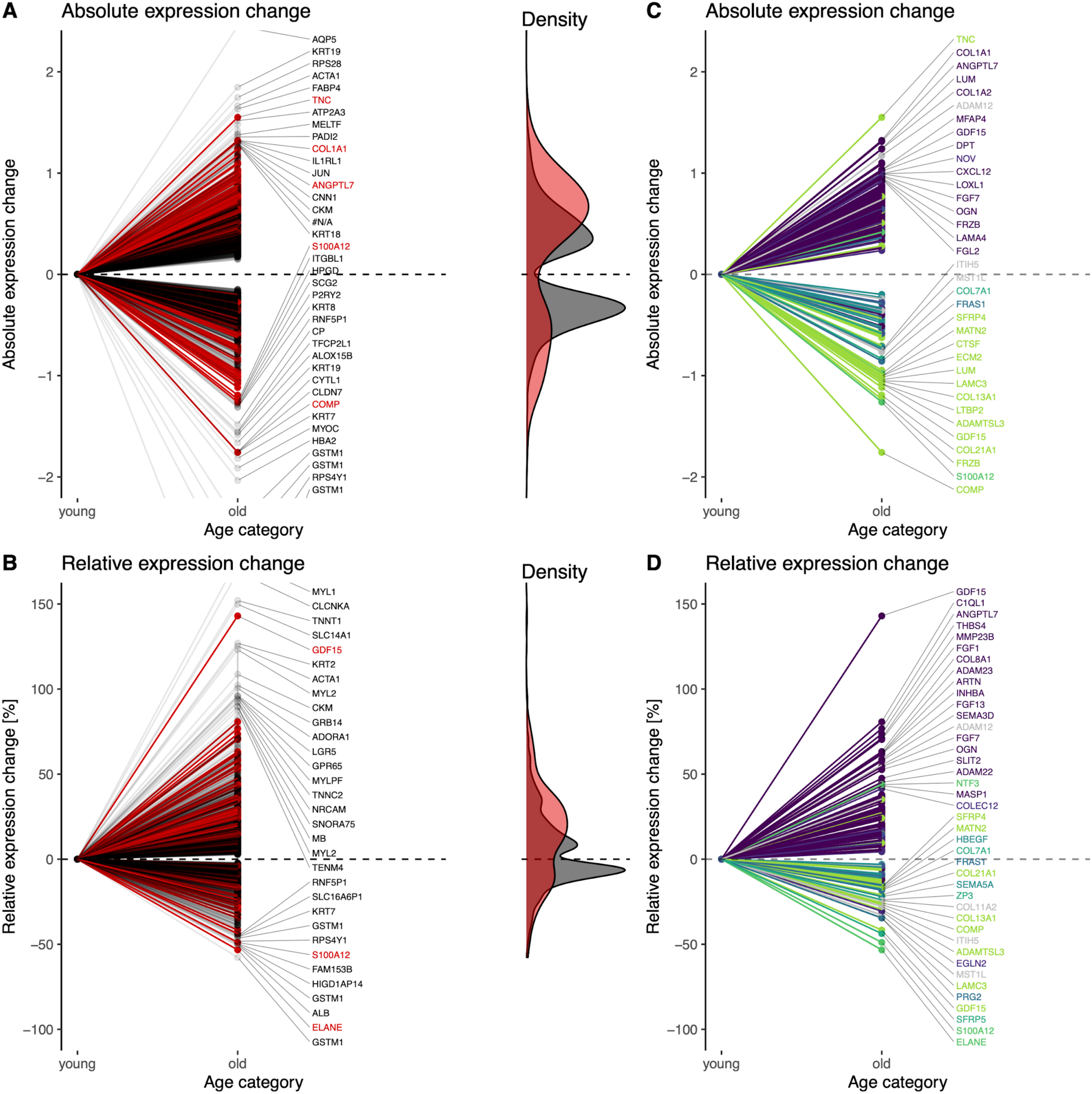
Matrisome genes affected by aging across human tissues. (A-B) Absolute (A) and relative (B) expression changes of all detected genes during aging are shown. Matrisome genes are displayed in red and non-matrisome genes in black. (C-D) Absolute (C) and relative (D) expression changes of only detected matrisome genes during aging are shown and are colored by tissue. For details, see Supplementary Table 3.

### Use of the young and aged matreotype to identify new pro-longevity compounds

The ultimate goal of our matreotype signature is for it to be used to identify new geroprotective compounds that modulate the matrisome, and thus, longevity. To first validate this approach, we used known pro-longevity compounds and identified those causing a youthful matreotype (*i.e.,* similar gene expression; Figure 4, Supplementary Tables 4, 5). We parsed the youthful matreotype signature into ‘downregulated matreotype genes that become upregulated during aging’ (Figure 4A) and ‘upregulated matreotype genes that become downregulated during aging’ (Figure 4B, Supplementary Table 4). We queried the CMap compound expression profiles and plotted the top 50 compounds that showed the gene expression signature as predicted by the youthful matreotype and called them ‘reversed aging signature’ compounds (Figure 4A, 4B). As a control, we also plotted the top 50 compounds that would enhance gene expression in the directionality of aging and called them ‘potentiated aging signature’ compounds (Figure 4A, 4B). Out of these total 200 compounds, 15 compounds were identified with both youthful or aged matreotype, so that we identified 185 unique compounds. Next, we searched the literature for reported lifespan increase in any organism upon treatment with any of these 185 compounds and found 24 of these compounds resulted in lifespan extension (Figure 4, Supplementary Table 5). To our surprise, these 24 compounds with reported lifespan increase did not cluster preferentially with the ‘reversed aging signature’ compounds as predicted but rather were almost equally distributed among all categories (Figure 4). This suggests that at least for matrisome genes, longevity might not be a simple reversion of gene expression associated with aging. Or it might be more complex given that each tissue has a unique matreotype (Supplementary Figures 5-7), such as that seen with GDF15 (Figure 3). Despite this unexpected finding, we note that our matreotype 100 gene compendium was able to identify 185 unique compounds, of which 13% have previously been reported to extend lifespan (Figure 4). This is an enrichment compared to the 5% of the 67 longevity-promoting compounds found in all 1309 CMap assessed small molecules (Figure 2A). Thus, independent of directionality, both youthful and age-related matreotype, *i.e.*, the matreotype signature itself predicts longevity-promoting drugs.

**Figure 4.**
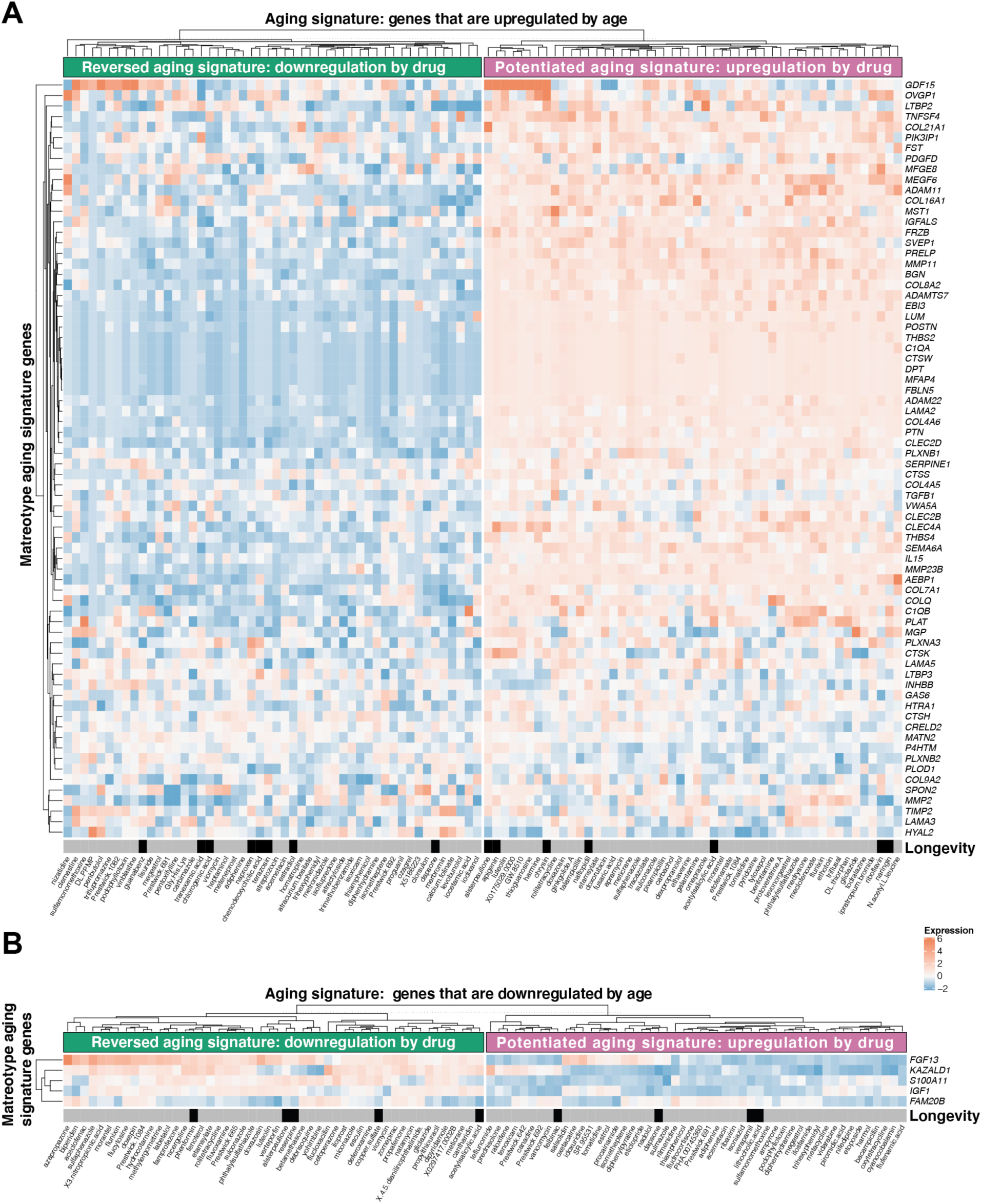
Matreotype signature gene expression correlating with CMap expression pattern. (A-B) Matreotype genes that are strongly upregulated (A) or downregulated (B) during aging are hierarchically clustered with gene expression patterns of small molecules on human cell lines from CMap library (Supplementary Table 4). Black rectangles indicate that the respective compound increases lifespan in any organism (Supplementary Table 5).

### Validating matreotype-predicted compounds with lifespan assays using *C. elegans*

To translate the *in-silico* analysis to an *in-vivo* functional relevance for healthy aging, lifespan assays in model organisms, such as *C. elegans* or mice are commonly used. The limitation of these lifelong assays, especially in mice, is that often one does not know if the optimal dose is applied until the end of the study three years later. To overcome this limitation, we developed an *in-vivo* screening assay measuring collagen biosynthesis. Similar to humans, collagen biosynthesis in *C. elegans* declines with age, and we recently discovered that many, if not all, longevity interventions prolong the expression of collagen genes in *C. elegans* (Ewald et al., 2015). This prolonged-expression of key collagen genes is required and sufficient for longevity (Ewald et al., 2015). Thus, we hypothesized that prolonged collagen expression, quantified on the transcriptional level by a collagen promoter-driven GFP, would constitute a useful surrogate marker to predict longevity (Figure 5A). Indeed, we found that the GFP intensity driven by collagen *col-144* promoter declined almost linearly within the first five days of adulthood (Figure 5B, Supplementary Table 6).

**Figure 5.**
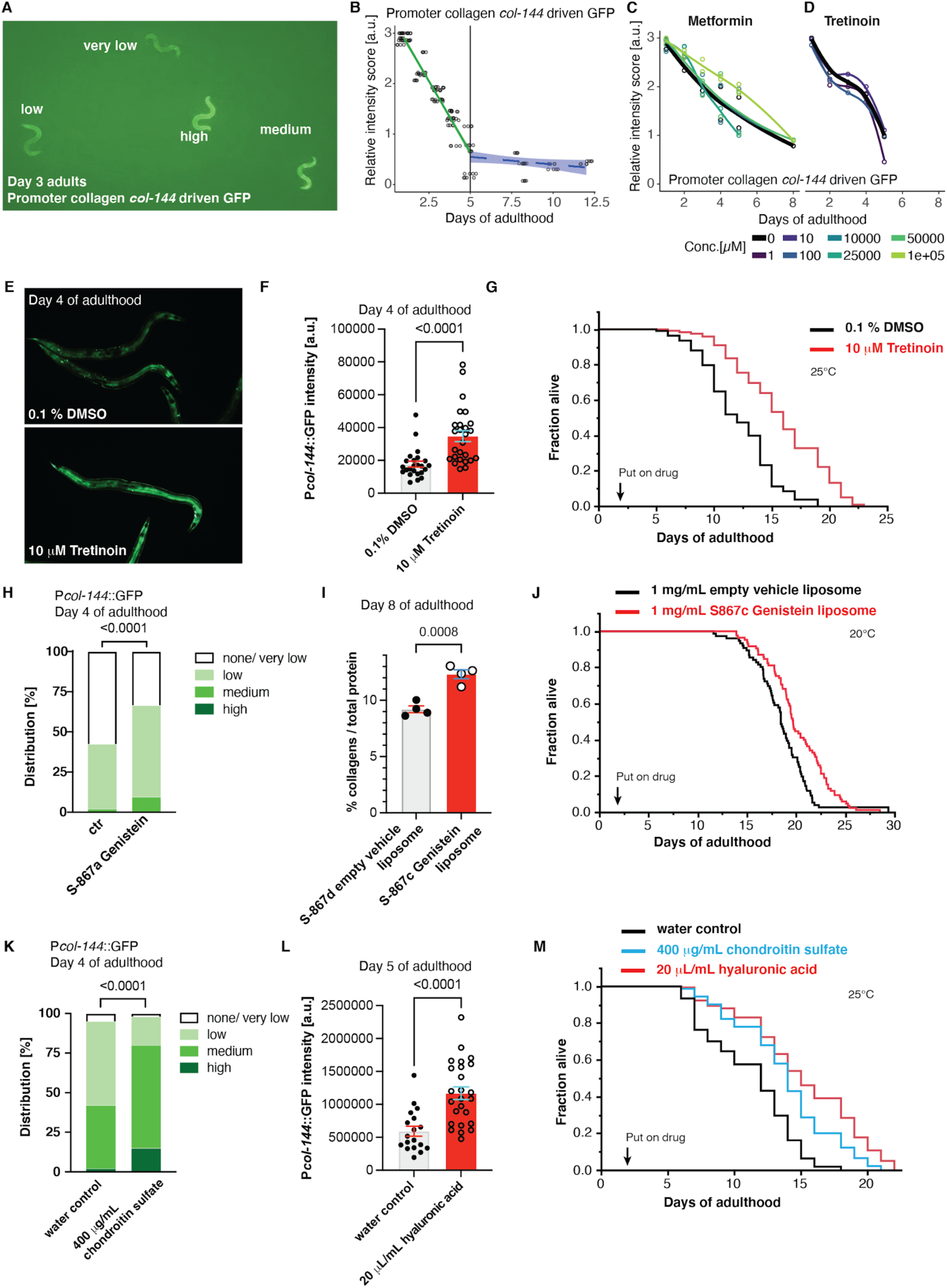
Validating compounds that were predicted to modulate the age-related matreotype for *in-vivo* collagen expression and lifespan. (A) Heterogeneity in decline of collagen *col-144* promoter driven GFP (LSD2002 [P*col-144*::GFP]) at day 3 of adulthood. GFP levels were categorized in 4 groups (none/very low, low, medium, high). (B) Approximately linear decline of P*col-144*::GFP expression during the first five days of adulthood. (C-D) Metformin (C) and tretinoin were observed to partially prolonged P*col-144*::GFP expression during aging in certain dosage regimes. (E-F) Representative image (E) and quantification (F) of P*col-144*::GFP expression at day-4 of adulthood upon tretinoin treatment from day-1 of adulthood. (G) Adulthood treatment (start at day-2 of adulthood) of tretinoin extended lifespan. (H) Adulthood treatment of genistein increased P*col-144*::GFP expression at day-4 of adulthood. (I) Liposomal-encapsulated genistein showed higher collagen over total protein levels at day-8 of adulthood. (J) Adulthood treatment (start at day-2 of adulthood) of liposomal-encapsulated genistein extended lifespan. (K) Adulthood treatment of chondroitin increased P*col-144*::GFP expression at day-4 of adulthood. (L) Adulthood treatment of hyaluronic acid increased P*col-144*::GFP expression at day-5 of adulthood. (M) Adulthood treatment (start at day-2 of adulthood) of chondroitin or hyaluronic acid extended lifespan. For raw data and statistics for (B-G, H, K-L), see Supplementary 6. For raw data, statistics, and additional trials (G, J, M), see Supplementary Table 7.

In our youthful matreotype-associated drugs, we have identified phenformin (Figure 4B, Supplementary Figure 9, Supplementary Table 2), a derivative of metformin. Both phenformin and metformin have been shown to increase *C. elegans* lifespan (Pryor and Cabreiro, 2015). Here, we found that treating almost adult (L4) *C. elegans* with metformin, prolonged collagen expression in a dose-dependent manner (Figure 5C; Supplementary Figure 10, Supplementary Table 6). This is consistent with a study showing metformin slows extracellular matrix morphological decline of the cuticle (Haes et al., 2014). This suggests that one unexplored aspect of the metformin’s mechanism-of-action might be via improved collagen homeostasis. Given this exciting finding, we decided to prioritize our investigations into drugs that will enhance collagen homeostasis in mammals but have not shown any pro-longevity phenotypes in any organisms. We, therefore, chose to test the retinoic acid receptor agonist tretinoin since tretinoin treatment prevents MMP upregulation and stimulates collagen synthesis in photo-aged skin (Griffiths et al., 1993). Our analysis showed that tretinoin had an enriched differential expression of matrisome genes (Supplementary Figure 9). Furthermore, tretinoin has been predicted to associate with a youthful expression pattern by the *in-silico* analysis of Janssens and colleagues but did not increase *C. elegans* lifespan at 50 μM (Janssens et al., 2019). With our reporter system, we found that treatment with 10 μM of tretinoin prolonged collagen expression and increased lifespan (Figure 5D-G; Supplementary Figure 10, Supplementary Table 6), confirming that tretinoin indeed promotes healthy aging.

Following the same rationale, in our drug hits that change matrisome expression, we identified genistein, an isoflavone (phytoestrogen) derived from soybeans to have a youthful matreotype profile (Supplementary Figure 9). Genistein has been predicted to associate with a youthful expression pattern by a previous *in-silico* analysis but failed to increase *C. elegans* lifespan at 50 μM (Janssens et al., 2019). By contrast, other groups reported a lifespan increase using 50 and 100 μM genistein (Lee et al., 2015). To reconcile this, we generated new 98% pure genistein and found prolonged collagen expression in aged *C. elegans* and a mild increase in lifespan (Figure 5H, Supplementary Figure 11, Supplementary Table 7). To optimize drug uptake, we encapsulated genistein extracts with liposomes. We found higher collagen protein content in aged *C. elegans* (day 8 of adulthood), enhanced oxidative stress resistance during older age (day 5 adults), and improved lifespan extension compared to empty vector liposomal control treatments (Figure 5I, 5J, Supplementary Figures 11-12, Supplementary Tables 7-8). A similar strategy was used to encapsulate royal jelly oil in a lecithin-based nanoemulsion for improved solubility on the NGM culturing plates to facilitate uptake by *C. elegans* (Supplementary Figure 11). These data indicate that optimizing drug delivery and dose is important for lifespan and healthspan benefits.

To expand our findings, we searched for compounds that could alter ECM composition and were not included in the CMap library. Besides collagens, glycoprotein and proteoglycans are the other two major components of ECMs across species (Naba et al., 2016; Teuscher et al., 2019a). We decided to investigate ECM precursors, such as glucosamine, chondroitin sulfate, and hyaluronic acid, which are part of the sugars added to these ECM proteins as potential mediators of the youthful matreotype. In humans, cohort studies of over 70-500 thousand participants who took glucosamine or chondroitin supplements showed a 15-18% and 22% reduction in total mortality, respectively (Bell et al., 2012a; Li et al., 2020). Here, we found that glucosamine treatment increased collagen expression during aging (Supplementary Figure 10, Supplementary Table 6). Previous studies also implicated glucosamine supplementation in lifespan extension of *C. elegans* and mice (Weimer et al., 2014). Using our collagen reporter screening system, we determined that 20 μL/ mL of hyaluronic acid and 400 μg/mL chondroitin sulfate prolonged collagen synthesis during aging and were sufficient to increase lifespan (Figure 5K-M; Supplementary Figures 10, 12, Supplementary Tables 6-8). Thus, our novel ECM-homeostasis reporter system is a powerful tool to identify drug-response doses that promote healthy aging.

## Discussion

Recent artificial intelligence, *in-silico*, and other computational approaches have been harnessed to predict beneficial and longevity-promoting effects of compounds, which were previously not considered to mediate effects on ECM gene expression (Bakula et al., 2018; Barardo et al., 2017b; Janssens et al., 2019; Moskalev et al., 2015; Vanhaelen et al., 2017). A major challenge lies in the validation of the health-promoting results of a novel compound. Here we demonstrated that a concise list of about 1000 matrisome or about 100 matreotype genes facilitates the identification of lifespan enhancing drugs. To establish a proof-of-concept for our matreotype approach, we developed a new non-invasive and *in-vivo* reporter system, which we used to validate known and novel geroprotective drugs. We used our system to determine the appropriate dose to unravel the compounds’ longevity potential indicated with our, and previous, *in-silico* approaches.

Established geroprotective drugs, such as metformin, rapamycin, resveratrol, and others, with known lifespan-extending effects, have previously been reported to alter the expression of ECM components (Figure 1). When we compared gene expression signatures, we found that almost ninety percent of the known longevity-promoting compounds in the CMap library showed changes in matrisome gene expression. To identify novel geroprotective drugs, we refined our approach by parsing ECM gene signatures resembling a young or aged matreotype to correlate with a given drug’s gene expression pattern. We found 185 candidate drugs, of which 24 showed lifespan increase in model organisms and 42 unique compounds had previously been predicted as potential geroprotectors (Supplementary Table 5).

The generally accepted assumption is that there is a drift in gene expression during aging, and that restoration of a younger gene expression pattern indicates rejuvenation of cells or tissues. This premise has been extensively used as a biomarker for the restoration of health in clinical trials (NCT02432287, NCT02953093), parsing healthy versus common aging cohorts (Zeng et al., 2020), reprogramming cells into a younger state (Lu et al., 2020), or in previous *in-silico* approaches (Tyshkovskiy et al., 2019). This premise also requires that longevity or rejuvenating interventions work through temporal scaling, a process that has been shown to be the case for lifespan extension and aging-associated gene expression in *C. elegans* (Stroustrup et al., 2016; Tarkhov et al., 2019) but not yet for mammals. We found that longevity-promoting drugs either increase or decrease matrisome gene expression (Figure 2). With a more refined approach using the youthful matreotype, we observed that both reversing or propagating the aging gene signatures could predict geroprotective drugs (Figure 4). One explanation could be that there is overlap in the gene expression signatures during aging and chronic diseases are similar (Fernandes et al., 2016; Wang et al., 2009a; Yang et al., 2015; Zeng et al., 2020).

Clearly, independent of directionality, the current defined age-related matreotype holds predictive power to identify new lifespan enhancing drugs. A shortcoming of our definition of the aging- and youth-associated expression signatures is that we do not take the context of the individual tissues into consideration. Unfortunately, insufficient studies are available to compile high-quality subsets for each tissue, which should be addressed in further experimental investigations. This is especially important in the case of collagen expression at an advanced age that can be both associated with improved tissue maintenance as observed in the skin and joints, while at the same time be implicated in fibrotic changes in the liver and kidney. For instance, downregulation of ECM in fat tissue but not in blood vessels is a key gene signature for healthy elderly individuals (Zeng et al., 2020). During aging, inflammation increases, which proceeds fibrosis (Wick et al., 2013) associated with 45% of deaths in the US alone (Wynn, 2008). On the other hand, collagen synthesis declines during aging and degradation or fragmentation by increased MMP activities, evident in aging skin (Fisher et al., 2009). Several drugs, including rapamycin (Chung et al., 2019), tretinoin (Griffiths et al., 1993), genistein (Polito et al., 2012), resveratrol (Lephart and Andrus, 2017), increase collagen synthesis in the skin. By contrast, resveratrol, rapamycin, and genistein suppress fibrotic collagen deposition in intestinal fibroblasts, kidney, or pulmonary fibrosis, respectively (Chen et al., 2012; Li et al., 2014; Matori et al., 2012). Thus, drugs might act as a geroprotector and increase lifespan by either inhibiting or enhancing the matrisome expression depending on pre-existing tissue damage or disease.

The model organism *C. elegans* does not show inflammation nor fibrosis during aging. Thus, collagen expression in *C. elegans* might reflect restoration or repair of the progressive decline of ECM homeostasis during aging analogous to human skin. This makes *C. elegans* the ideal readout for any age-dependent changes of matrisome genes observed from human or mammalian omics approaches. Based on this, we established an age-dependent collagen homeostasis read-out as a predictive marker for longevity.

In previous work, more than 100’000 compounds have been screened, and about 100 compounds have been identified to increase *C. elegans* lifespan (Supplementary Table 5) (Kim et al., 2019; Lucanic et al., 2013; Petrascheck et al., 2007; Ye et al., 2014). A practical limitation in verifying drug candidates is the unknown dosage to be used. Usually, one dose is chosen for all compound treatments for *C. elegans* lifespan screening assays, potentially leading to many false-negative results and limiting its interpretation. Dose-response curves are not linear, but often J- or U-shaped, whereby in general, high doses are toxic and low doses lead to hormetic responses increasing lifespan (Ristow and Schmeisser, 2014). We showed that two previously predicted but regarded as false-positive compounds - tretinoin and genistein-robustly increase lifespan when assayed at the appropriate dosage. Furthermore, optimization for the route of uptake improves robustness, promoting healthy aging. Thus, our novel reporter system serves as a predictive tool to identify the appropriate dosage for lifespan assays.

It is striking that longevity-promoting compounds identified in model organisms showed enriched matrisome gene expression signature in human cells treated with these compounds (Supplementary Figures 2-4). The proposed mechanisms for longevity-compounds discovered with *C. elegans*, mostly work through intercellular communication (Petrascheck et al., 2007; Ye et al., 2014). Pathway analysis showed an enrichment for chondroitin and heparan sulfate biogenesis and TGFβ pathway as predicted drug-protein targets (Liu et al., 2016). On the other hand, the outcome of previous *in-silico* approaches using youthful gene expression determined from GTEx data to query CMap identified HSP90 chaperone network (Dönertaş et al., 2019; Fuentealba et al., 2019; Janssens et al., 2019) and protein homeostasis (Komljenovic et al., 2019) as healthy aging promoting interventions. HSP90 is found in the extracellular space binding fibronectin and chaperones other ECM proteins (Baker-Williams et al., 2019; Hunter et al., 2014). Pharmacological inhibition of HSP90 alters ECM signaling (Chaturvedi et al., 2011). Proper ECM protein homeostasis is essential to ensure intercellular communication, a hallmark lost during aging (Ewald, 2019; López-Otín et al., 2013). This raises the question whether geroprotective drugs improve ECM homeostasis. There is tantalizing evidence with established longevity-promoting medications, such as rapamycin, resveratrol, metformin, and others (Figure 1B), and we provided experimental evidence for this with tretinoin, genistein, glucosamine, chondroitin, and hyaluronic acid (Figure 5). Consistent with this is that glucosamine and chondroitin stimulate collagen synthesis *in vitro* and *ex vivo* of elderly human skin in a clinical trial (Gueniche and Castiel-Higounenc, 2017; Lippiello, 2007), extracellular matrix component hyaluronic acid treatments stimulate collagen synthesis in human photo-aged skin (Wang et al., 2007), and in mice (Fan et al., 2019), and topical application of rapamycin restores collagen VII levels in a clinical trial (NCT03103893) (Chung et al., 2019). ECM homeostasis might remodel or prevent glycation and crosslinking of collagens (Ewald, 2019). There are currently 27 clinical trials addressing ECM stiffness and its role in diseases by investigating eleven different molecular targets (Lampi and Reinhart-King, 2018). Furthermore, different matreotypes might be valuable prognostic factors or biomarkers (Ewald, 2019). For instance, ECM is one of the strongest associated aging-protein signatures in blood plasma or urine proteomics of healthy older adults (Nkuipou-Kenfack et al., 2015; Sathyan et al., 2020). Thus, defining matreotypes related to healthy aging or age-related chronic diseases might be a strategy for personalized medicine approaches.

In summary, we demonstrated that prolonged ECM homeostasis is a biomarker for *C. elegans* longevity and harnessed this to establish a novel *in-vivo* assay. We provided evidence that gene expression patterns of human cells treated with known geroprotective drugs alter ECM genes, and developed an age-stratified matreotype. We then used this matreotype to identify novel geroprotective compounds based upon their transcriptomes. Our method highlights a previous unused potential of ECM reprogramming as a means to identify and validate novel compounds, licensed drugs, natural compounds, and supplements that potentially retard or prevent age-related pathologies. Understanding pharmacological reprogramming of extracellular environments may provide a new platform to discover previously unidentified therapeutic avenues and holds significant translational value for disease diagnostics.

## Materials and Methods

### Strains

*Caenorhabditis elegans* strains were maintained on NGM plates and OP50 *Escherichia coli* bacteria. The wild-type strain was N2 Bristol. Mutant strains used are described at www.wormbase.org: TJ1060: *spe-9(hc88) I; rrf-3(b26) II.* LSD2002 was generated by integrating P*col-144*::GFP transgene (a generous gift from Yelena Budovskaya and Stuart Kim (Budovskaya et al., 2008)) into N2, outcrossing eight times, and crossing to TJ1060, resulting *spe-9(hc88) I; xchIs001* [P*col-144*:: GFP; *pha-1*(+)] X genotype.

### Data analysis

Data analysis was performed utilizing the dplyr (Hadley Wickham, Romain François, Lionel Henry and Kirill Müller (2020). dplyr: A Grammar of Data Manipulation. R package version 1.0.0.) and purrr (Lionel Henry and Hadley Wickham (2020). purrr: Functional Programming Tools. R package version 0.3.4.) R packages. Data visualization was generated using ggplot2 (H. Wickham. ggplot2: Elegant Graphics for Data Analysis. Springer-Verlag New York, 2016.), ComplexHeatmap (Gu, Z. (2016) Complex heatmaps reveal patterns and correlations in multidimensional genomic data. Bioinformatics.), and ggpubr (Alboukadel Kassambara (2020). ggpubr: ‘ggplot2’ Based Publication Ready Plots. R package version 0.4.0.999.).

### Data origin

Human matrisome (http://matrisomeproject.mit.edu, (Naba et al., 2016)), GTEx data (Consortium, 2013), age change (Janssens et al., 2019). CMap data (Lamb et al., 2006), lifespan information were obtained from GenAge (Magalhães and Toussaint, 2004), Geroprotectors (Moskalev et al., 2015), and through a comprehensive literature review.

### Literature search

Compounds that extended the mean lifespan of the organism by more than 5%, according to the DrugAge (Barardo et al., 2017a) and Geroprotector (Moskalev et al., 2015) databases, were selected for further investigation. The abstracts of studies associated with lifespan extension were filtered for the occurrence of at least one of the following matrisome keywords: collagen, ECM, extracellular, matrix, proteoglycan, hyaluronic, hyaluronan, TGF, integrin, TGFbeta.

### Aging matreotype definition

To define the human aging matreotype, we performed a literature search and extracted the age-association of all genes involved in forming the human matrisome. We aggregated information stemming from multiple different data types ranging from age-dependent differential expression (Komljenovic et al., 2019; Yang et al., 2020), age-association (Dönertaş et al., 2018), age-correlation of expression (Yang et al., 2015), gene products directly linked to aging or linked to known aging regulators (Wang et al., 2009b), manually curated age-related genes or regulating genes related to aging (Magalhães and Toussaint, 2004), a mammalian meta-analysis of age-regulation (Magalhães et al., 2009), transcriptome biomarkers of healthy aging (Sood et al., 2015), age-associated DNA methylation (Bacalini et al., 2015; Bell et al., 2012b). A large part of aging datasets were obtained from a large-scale meta-analysis conducted by (Blankenburg et al., 2018). If the datasets have not yet been subjected to a significance cutoff, we applied multiple testing corrected (Benjamini-Hochberg) threshold of 0.05 to each dataset if applicable. Studies analyzing individual tissues were treated as separate sources. To define the aging matrisome, we acquired data from at least three sources implicating the gene in the aging process. Studies that offer directionality were further utilized to determine matrisome genes that were upregulated or downregulated with age using the same thresholds.

### Collagen promoter-driven GFP scoring

The strain LSD2002 P*col-144::GFP* was used to assess age-related decline in collagen-144 expression. Animals were age-synchronised by bleaching, and made infertile by culturing at 25℃. NGM culturing plates were prepared with tretinoin (Sigma PHR1187), hyaluronic acid (Sigma H5388), chondroitin sulfate (Sigma 27042), metformin (Sigma D150959), glucosamine (Sigma G4875), and genistein (Changzhou Longterm Biotechnology Co.,Ltd). Distribution scoring is based on intensity observed visual inspection, categorizing to none or very low, low-, medium-, or high intensity. For some experiments, animals were mounted onto 2% agar pads and pictures were taken with an upright fluorescent microscope (Tritech Research, model: <u>BX-51-F</u>). To separate the GFP signal from the autofluorescence of the gut, we used the microscope, settings and triple-band filterset as described by Teuscher (Teuscher and Ewald, 2018). The total intensity per animal, intensity [a.u.], is calculated from fluorescence images using FIJI.

### Manual lifespan measurements

Manual scoring of lifespan as previously described by Ewald *et. al.* 2016 (Ewald et al., 2016). In brief, about 100 day-2 adult *C. elegans* were picked to NGM plates containing the solvent either water or 0.1% dimethyl sulfoxide (DMSO) alone as control or tretinoin (Sigma PHR1187), hyaluronic acid (Sigma H5388), chondroitin sulfate (Sigma 27042). Animals were classified as dead if they failed to respond to prodding. Exploded, bagged, burrowed, or animals that left the agar were excluded from the statistics. The estimates of survival functions were calculated using the product-limit (Kaplan-Meier) method. The log-rank (Mantel-Cox) method was used to test the null hypothesis and calculate *P* values (JMP software v.14.1.0.).

### Automated survival assays using the lifespan machine

Automated survival analysis was conducted using the lifespan machine described by Stroustrup and colleagues (Stroustrup et al., 2013). Approximately 500 L4 animals were resuspended in M9 and transferred to NGM plates containing 50 µM 5-Fluoro-2’deoxyuridine (FUdR) seeded either with OP50 bacteria, or with RNAi bacteria supplemented with 100 μg/ml carbenicillin, or with heat-killed OP50 bacteria, or with UV-inactivated *E. coli* strain NEC937 B (OP50 uvrA; KanR) containing 100 μg/ml carbenicillin. Animals were kept at 20°C until measurement. Tight-fitting Petri dishes (BD Falcon Petri Dishes, 50 x 9 mm) were used for lifespan experiments. Tight-fitting plates were dried without lids in a laminar flow hood for 40 minutes before starting the experiment. Air-cooled Epson V800 scanners were utilized for all experiments operating at a scanning frequency of one scan per 10 – 30 minutes. Temperature probes (Thermoworks, Utah, U.S.) were used to monitor the temperature on the scanner flatbed and maintain 20°C. Animals that left the imaging area during the experiment were censored. Population survival was determined using the statistical software R (Ihaka and Gentleman, 2012) with the survival (Therneau and Grambsch, 2000) and survminer (https://rpkgs.datanovia.com/survminer/) packages. Lifespans were calculated from the L4 stage (= day 0).

### Compound preparation for lifespan and oxidative stress assays

Compounds are received freeze-dried, except for the liposomes, which were acquired in 100% saturated suspension. All compounds were blinded with a serial number. Compounds and control solvents are administered to *C. elegans* by mixing it in the Nematode Growth Medium (NGM) immediately before pouring the plates. Compound stock solutions were made by dissolving 100 mg/mL in their solvent, 100% DMSO for genistein, water for the royal jelly oil in lecithin-based nanoemulsion. The royal jelly oil was prepared from dispersing royal jelly powder in oil (mygliol) but it was not soluble in the NGN agar *C. elegans* culturing plates. In a second step, we encapsulated the royal jelly oil by homogenizing the oil with lecithin. The liposomal genistein and empty liposomes were suspended in water. These stocks were consequently used to make dilution series. The final concentration of DMSO on the lifespan plates did not exceed 0.2%. The *C. elegans* strain TJ1060 was age-synchronized by extracting the eggs with bleach and were made infertile by culturing at 25°C from egg to day-1 of adulthood. On day-2-of adulthood, 30-40 animals were placed per 6 cm plates, four plates for each compound. Subsequently, the plates are loaded onto the scanners, kept in a controlled environment at 20°C. Every scanner includes at least four control plates. For the manual lifespan at 25°C, three plates were used per compound, and death events were counted once per day.

### Oxidative stress assays

Oxidative stress assay was modified from Ewald *et al.,* 2017 (Ewald et al., 2017). *C. elegans* of the L1 or day-1-adult stage were shifted on compound-containing or control plates washed off at the indicated time point, incubated with 14 mM sodium arsenite (Honeywell International 35000) in U-Shaped 96 well plates, and put into the wMicroTracker (MTK100) for movement scoring. For statistical analysis, the area under the curve was measured, and the mean for each run was calculated. Statistical analysis was performed by using a paired sample *t*-test.

### Quantifying total collagen over protein content

Collagen levels were determined by hydroxyproline content as described in Teuscher *et al*., 2019 (Teuscher et al., 2019b). In brief, about 10 000 TJ1060 *C. elegans* eggs were placed at 25°C until day-1 of adulthood and then transferred on plates containing the compounds at 20°C. Day-8-adult animals were harvested for the collagen and protein assays.

## Author contributions

All authors participated in analyzing and interpreting the data. CYE and CS designed the experiments. CS established and performed *in-silico* analysis. AD and PM translated compound names for list comparison and automated analysis. EJ, CS, and CYE performed lifespan assays. EJ and CYE performed oxidative stress assays. EJ and SXL performed collagen expression assays. FW and FZ prepared and encapsulated compounds. CYE wrote the manuscript in consultation with the other authors.

## Author Information

The authors have no competing interests to declare. Correspondence should be addressed to C. Y. E.

## Acknowledgement

We thank Anne Häfke for help with analysing GFP expression, Katharina Tarnutzer for help with the hydroxyproline collagen measurements, Alina Teuscher for LSD2002 strain, and Sarah Jayne Mitchell and Micheal Robert Mac Arthur for critical reading commenting on the manuscript. Some strains were provided by the CGC, which is funded by NIH Office of Research Infrastructure Programs (P40 OD010440). EJ was funded by a grant of the Mibelle Group Biochemistry, Mibelle AG, Switzerland. Funding from the Swiss National Science Foundation PP00P3_163898 to CYE and CS.

---

**Supplementary Figure 1.**
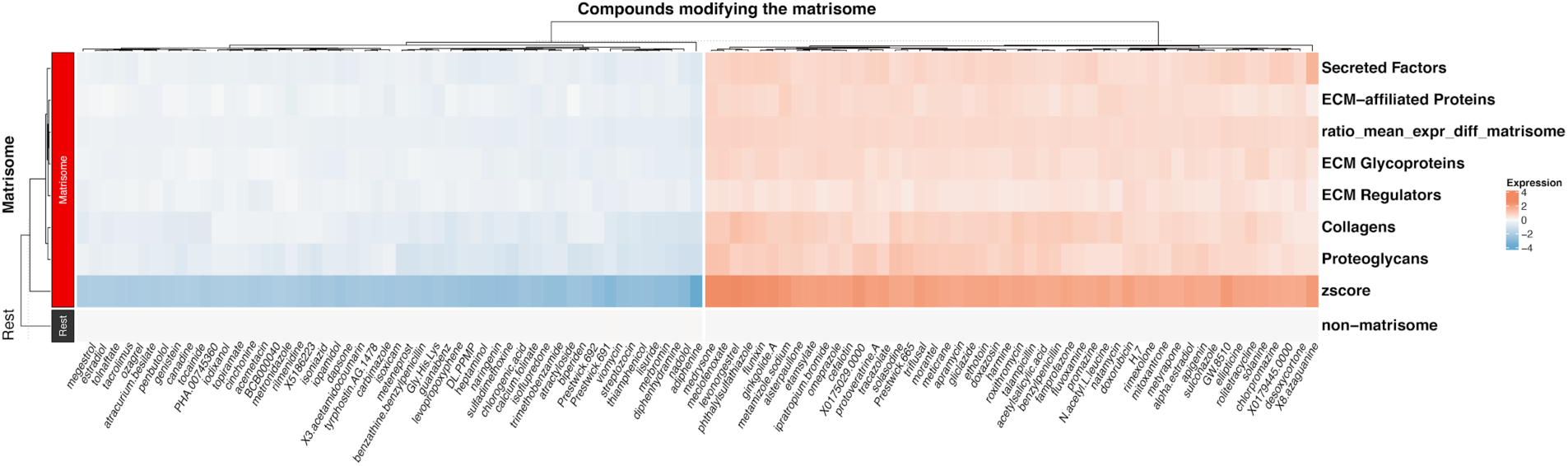
Matrisome categories being preferentially targeted by each compound. Compounds affecting matrisome gene expression are shown as the normalized expression for the entire matrisome (z-score), the matrisome subcategories (Collagens, ECM-affiliated Proteins, ECM Glycoproteins, ECM Regulators, Proteoglycans, and Secreted factors), the mean expression of the non-matrisome genes and the difference between the overall mean expression of the matrisome and non-matrisome (rest).

**Supplementary Figure 2.**
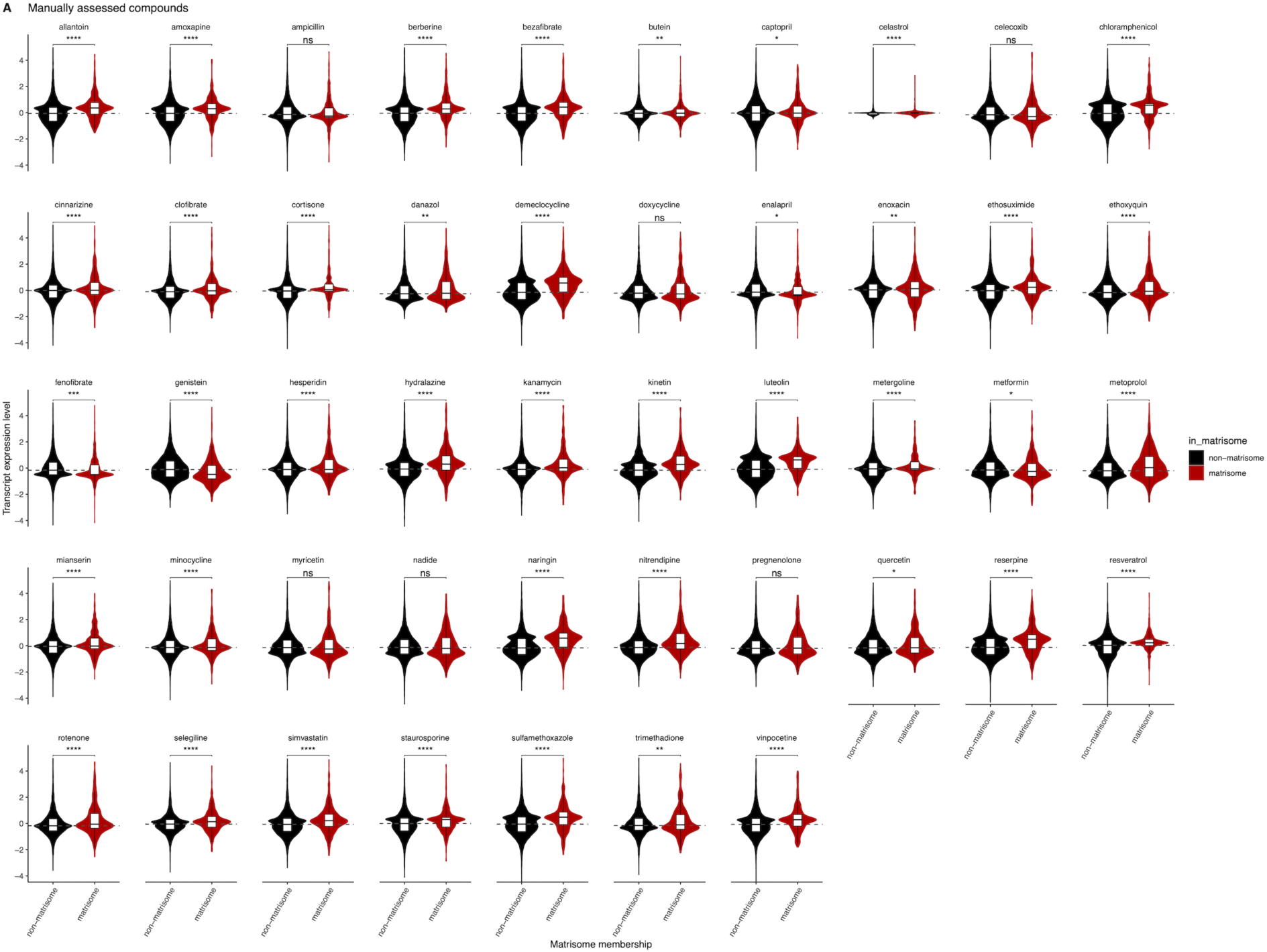
Overall matrisome regulation by compounds that were selected for experimental validation. The overall expression regulation of matrisome genes was assessed for 47 compounds using the CMap library dataset. The distribution of matrisome and non-matrisome genes is shown as both a box- and violin plot and the P-value of the Wilcox test is shown (ns: p > 0.05, *: p <= 0.05, **: p <= 0.01, ***: p <= 0.001, ****: p <= 0.0001).

**Supplementary Figure 3.**
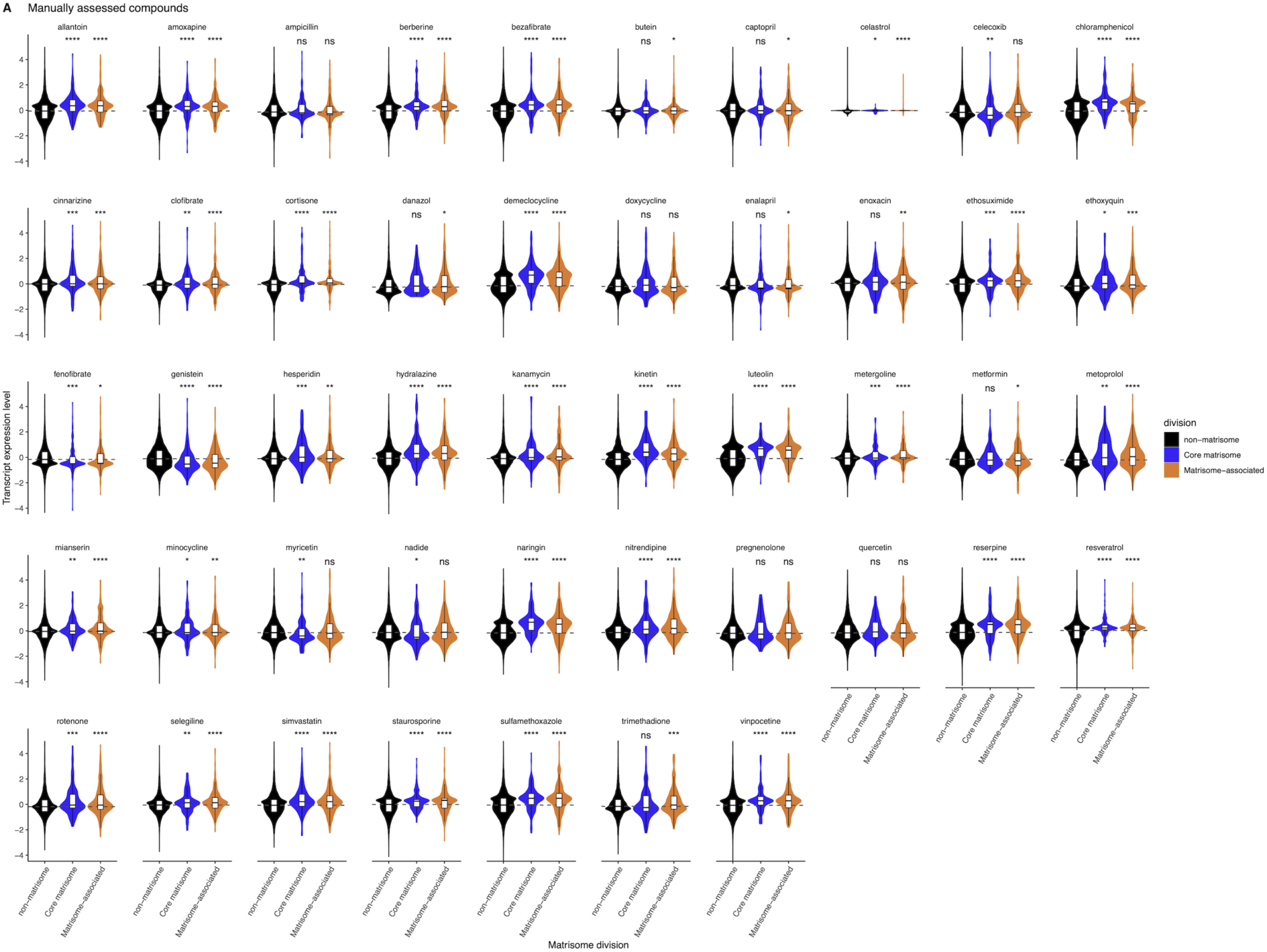
Specific regulation of matrisome divisions by compounds that were selected for experimental validation. The overall expression regulation of matrisome genes was assessed for 47 compounds using the CMap library dataset. The distribution of matrisome and non-matrisome genes is shown as both a box- and violin plot and the P-value of the Wilcox test is shown (ns: p > 0.05, *: p <= 0.05, **: p <= 0.01, ***: p <= 0.001, ****: p <= 0.0001).

**Supplementary Figure 4.**
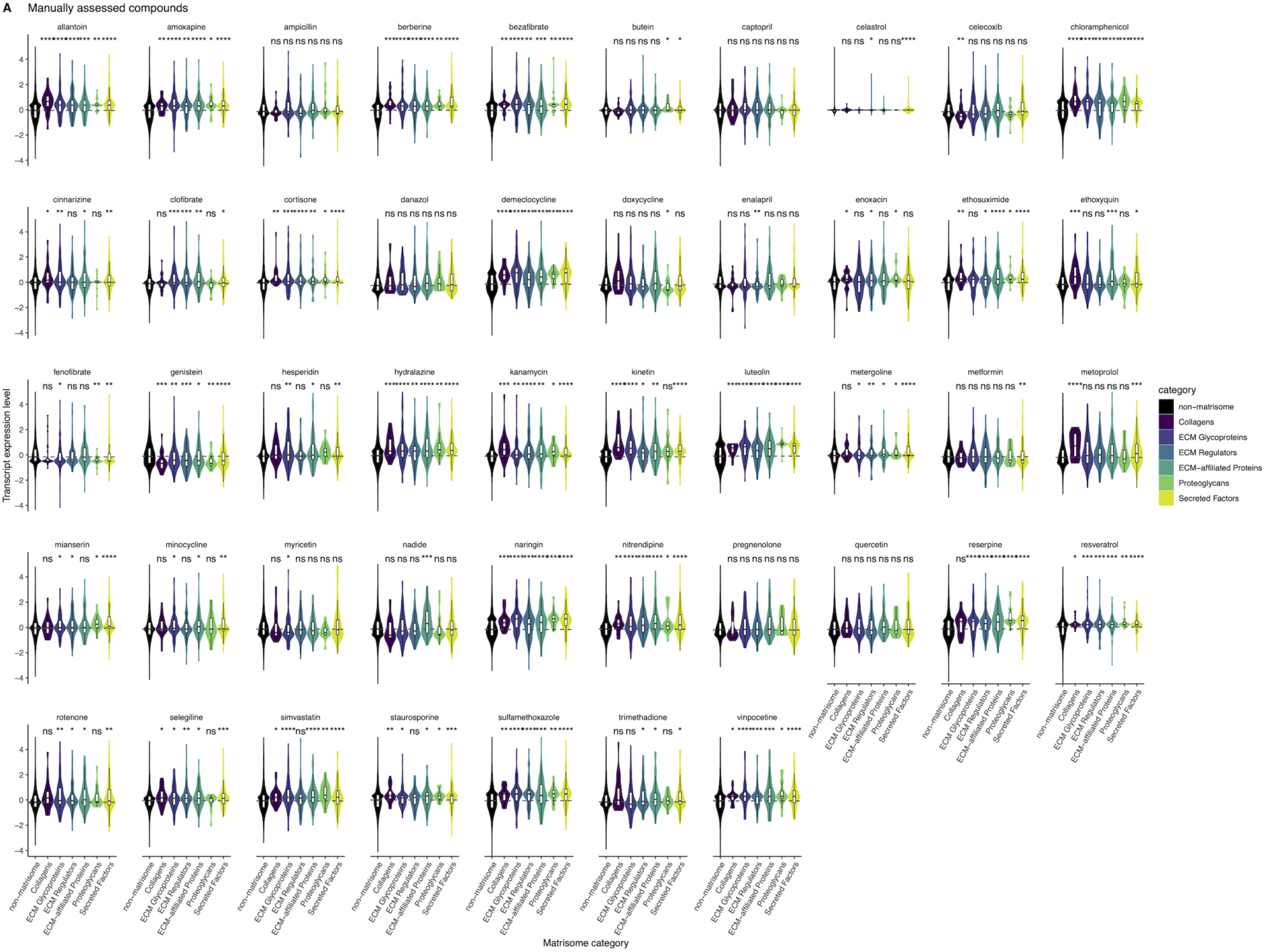
Specific regulation of matrisome categories by compounds that were selected for experimental validation. The overall expression regulation of matrisome genes was assessed for 47 compounds using the CMap library dataset. The distribution of matrisome and non-matrisome genes is shown as both a box- and violin plot and the P-value of the Wilcox test is shown (ns: p > 0.05, *: p <= 0.05, **: p <= 0.01, ***: p <= 0.001, ****: p <= 0.0001).

**Supplementary Figure 5.**
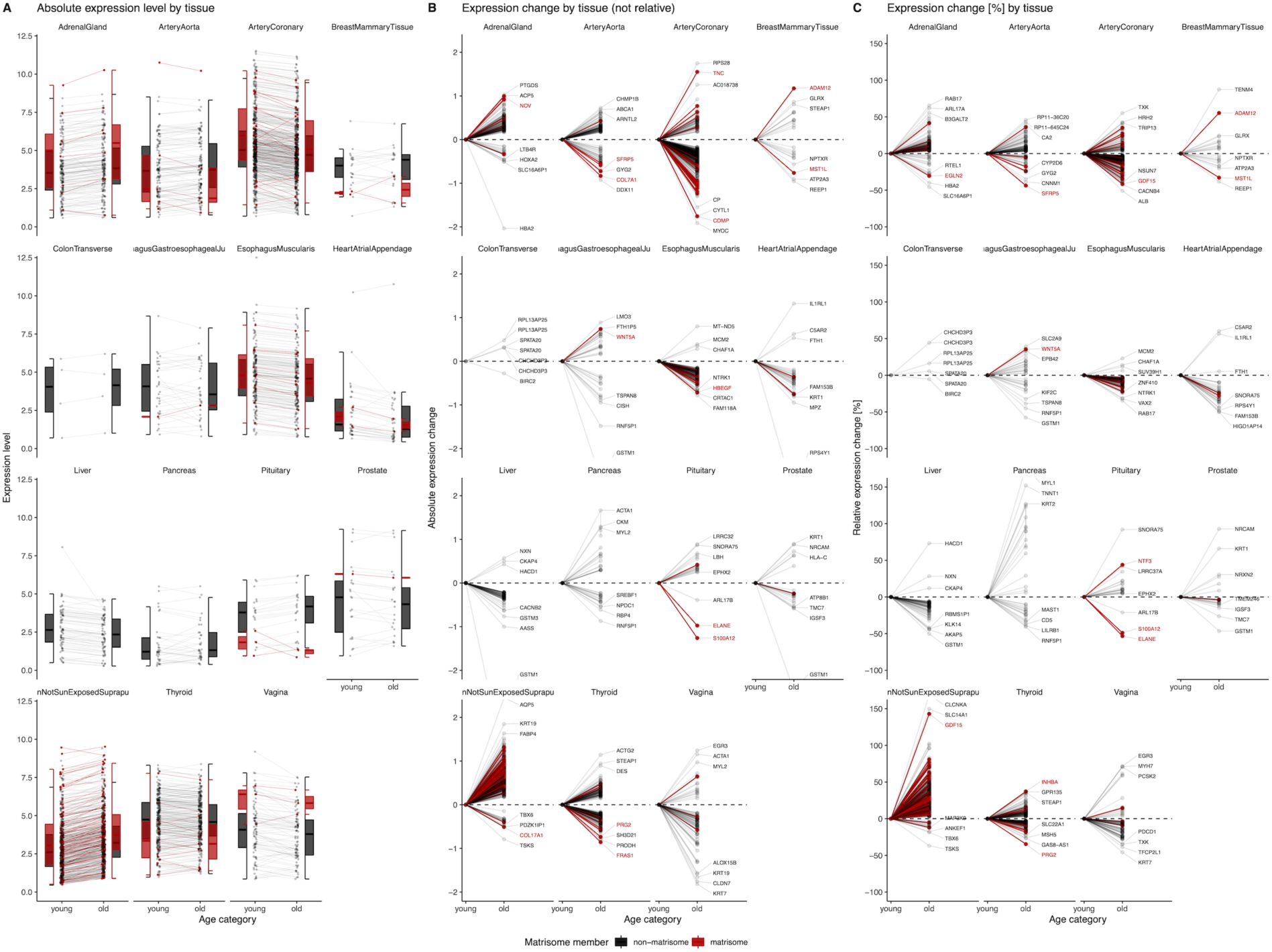
Matrisome gene expression across tissues. The gene expression signatures of old tissues were compared to the expression signature of the corresponding young tissues while highlighting the matrisome genes. Absolute expression (A), the change of absolute expression with age (B), and the relative expression change with age (C) are shown. The most differentially regulated genes are labelled. Each tissue is shown in its own panel, the matrisome genes in red, and the non-matrisome genes in black.

**Supplementary Figure 6.**
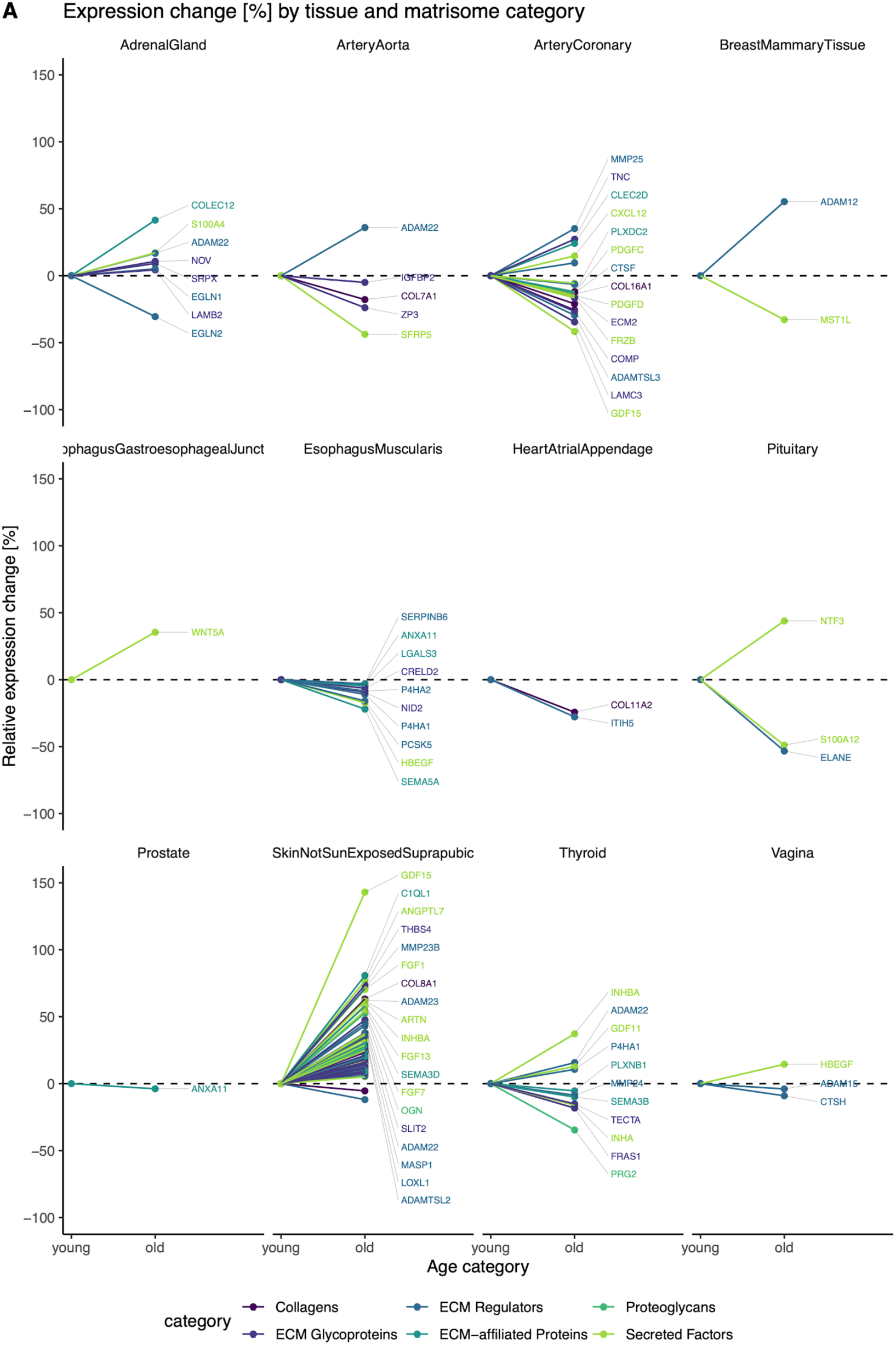
Individual involvement of matrisome categories in expression signatures accompanying tissue aging. The relative change in expression with age is displayed for all matrisome genes. Each gene is colored by its matrisome category.

**Supplementary Figure 7.**
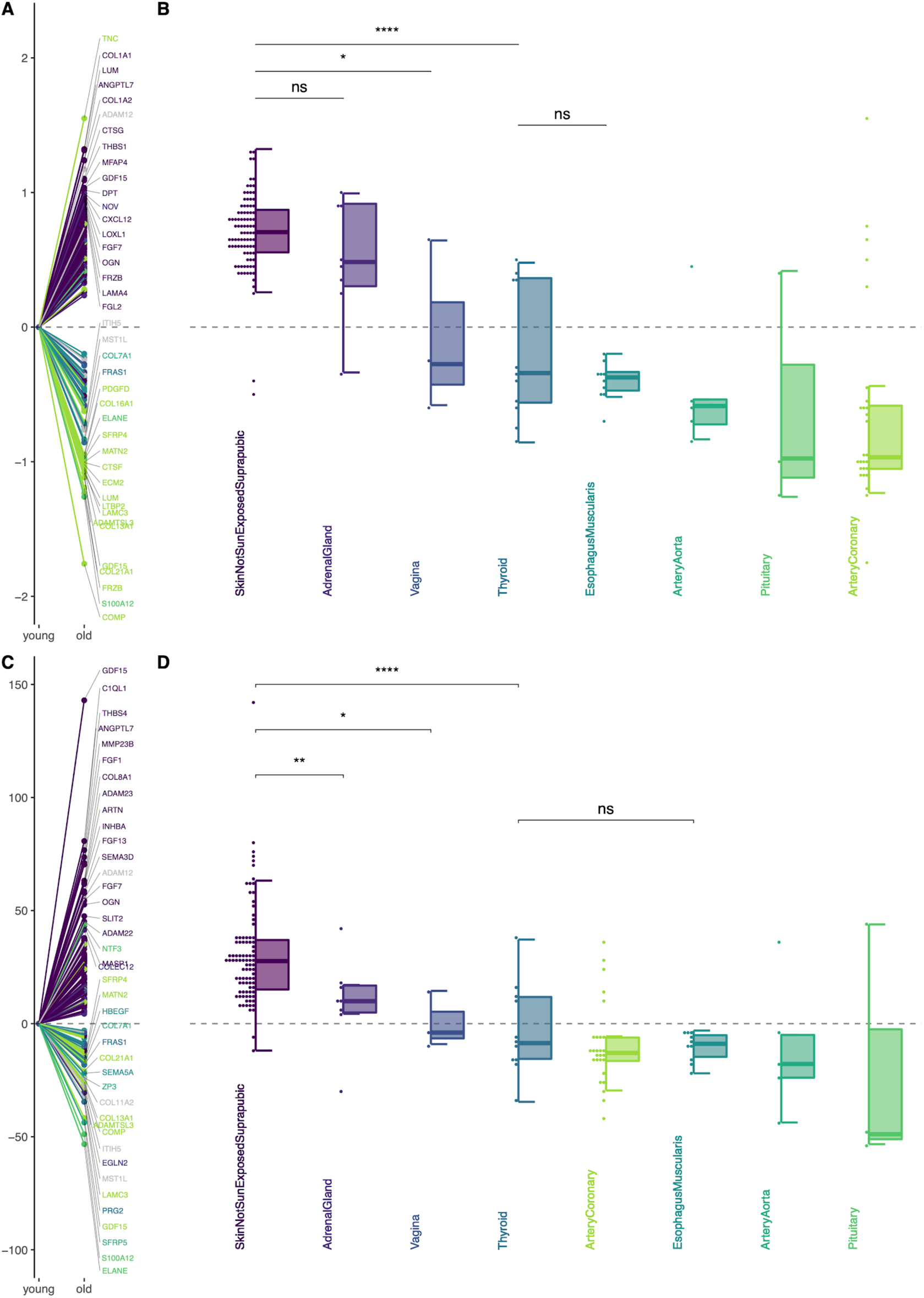
Individual involvement of matrisome genes in expression signatures accompanying tissue aging. The absolute (A, B) and relative [%] (C, D) expression change of matrisome genes in tissue aging is displayed and colored by tissue. Genes are compared relative to each other in (A) and (C) while the overall change on the level of each tissue is compared in (B) and (D). The P-value of the Wilcox test comparing selected tissues is shown (ns: p > 0.05, *: p <= 0.05, **: p <= 0.01, ***: p <= 0.001, ****: p <= 0.0001)

**Supplementary Figure 8.**
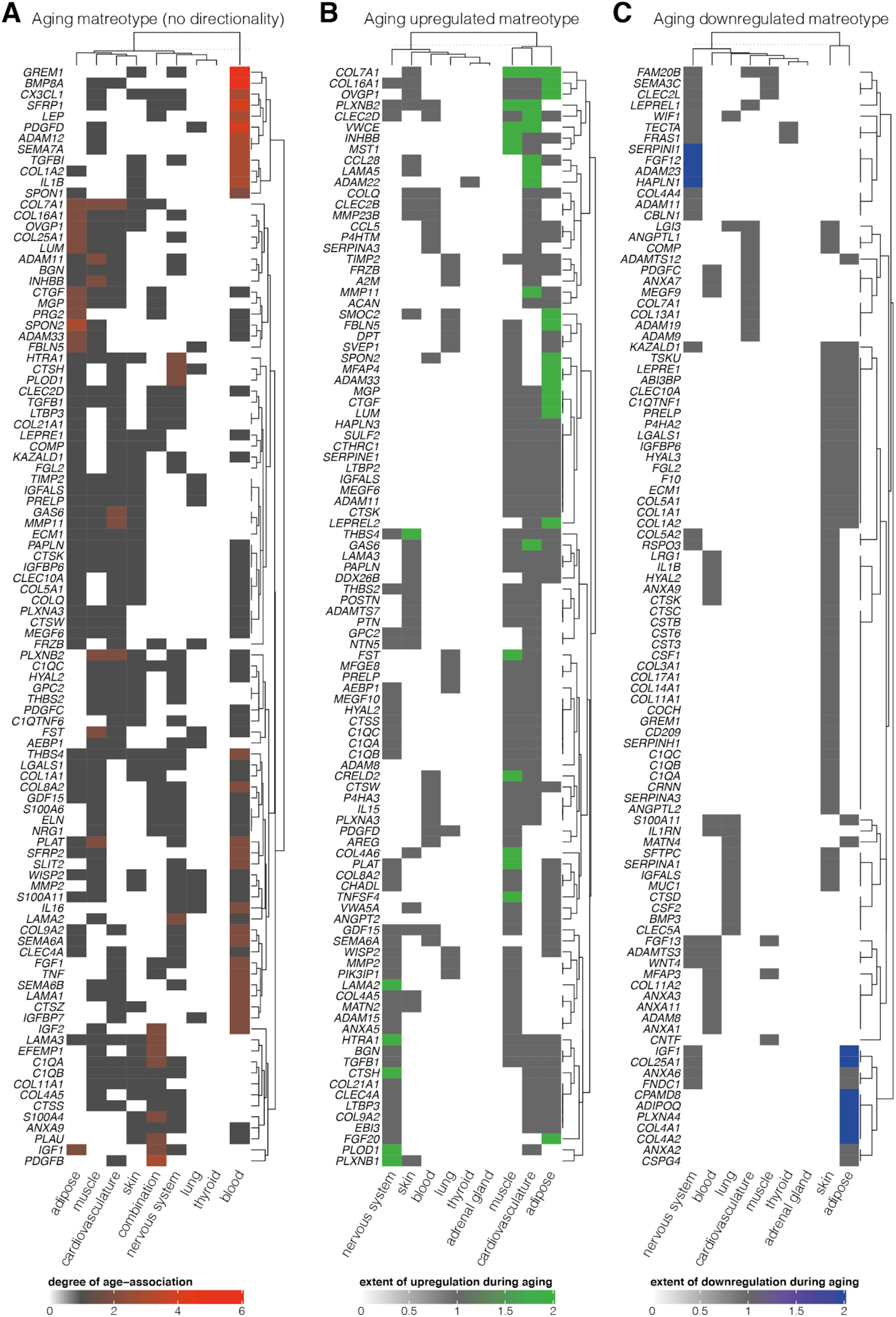
Genetic fingerprint of the aging human matreotype. Meta analysis of 48 human aging datasets to characterize matrisome aging on a single-gene level. In (A), studies were gathered which implicated matrisome genes in human aging without determining the regulation of the specific genes. A different perspective is provided by studies which assess the age-specific regulation change for each gene that are shown in (B) as up-regulation) and (C) asdown-regulation. The rows of the heatmap refer to the most age-regulated matrisome genes with the tissues shown as columns in which a significant age-associated link has been observed for each gene. The color of each heatmap cell corresponds to the number of studies in support of this age-association. Both genes and tissues are clustered hierarchically to identify shared aging patterns.

**Supplementary Figure 9.**
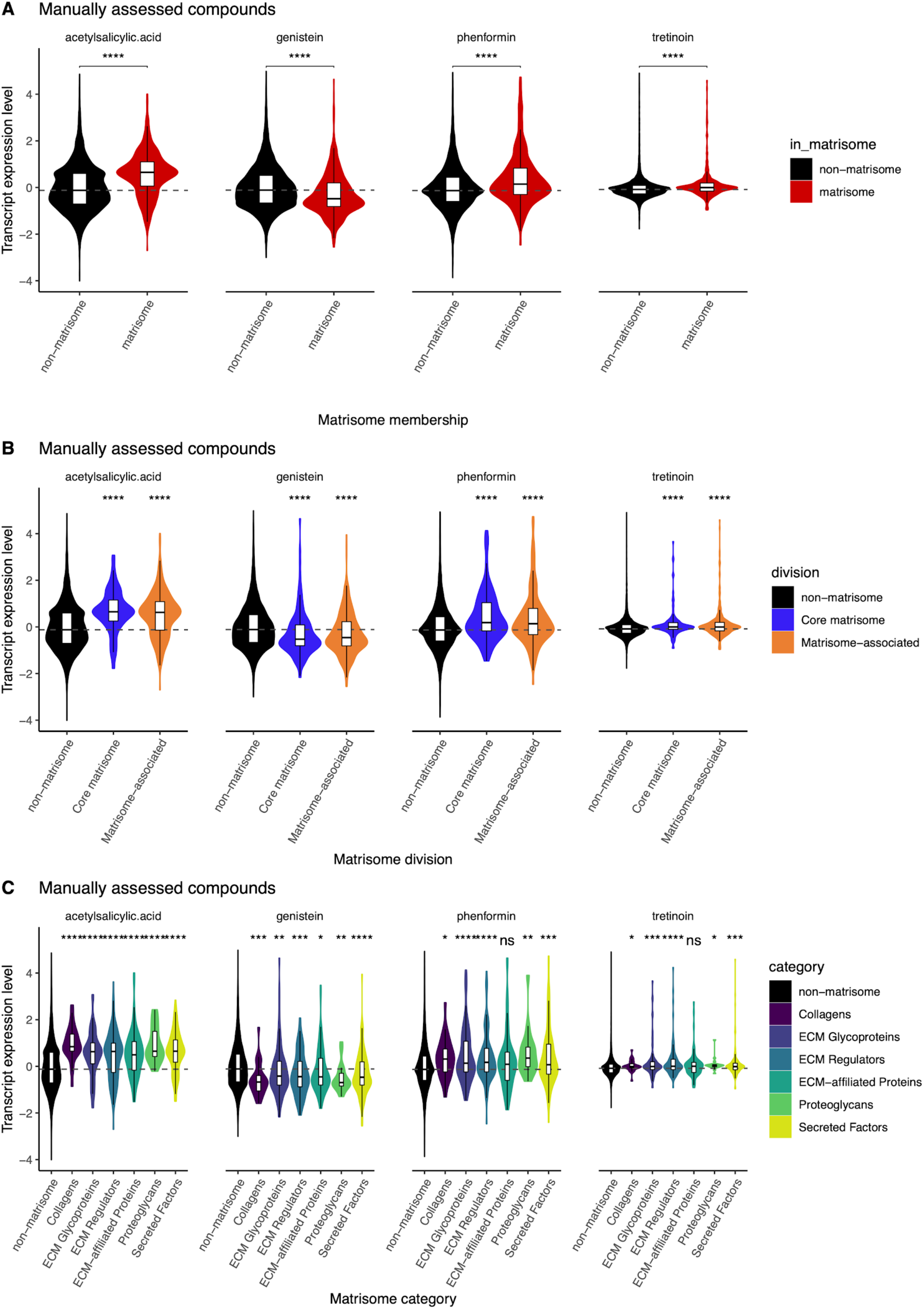
Regulation of matrisome gene expression of compounds experimentally assessed for collagen gene expression and lifespan extension in *C. elegans*. The expression regulation of the matrisome as a whole (A), the individual matrisome divisions (B) and categories (C) by acetylsalicylic acid, genistein, phenformin, and tretinoin were analyzed using the CMap library dataset. The distribution of matrisome and non-matrisome genes is shown as both a box- and violin plot and the P-value of the Wilcox test of each gene group compared to the non-matrisome subset is shown (ns: p > 0.05, *: p <= 0.05, **: p <= 0.01, ***: p <= 0.001, ****: p <= 0.0001)

**Supplementary Figure 10.**
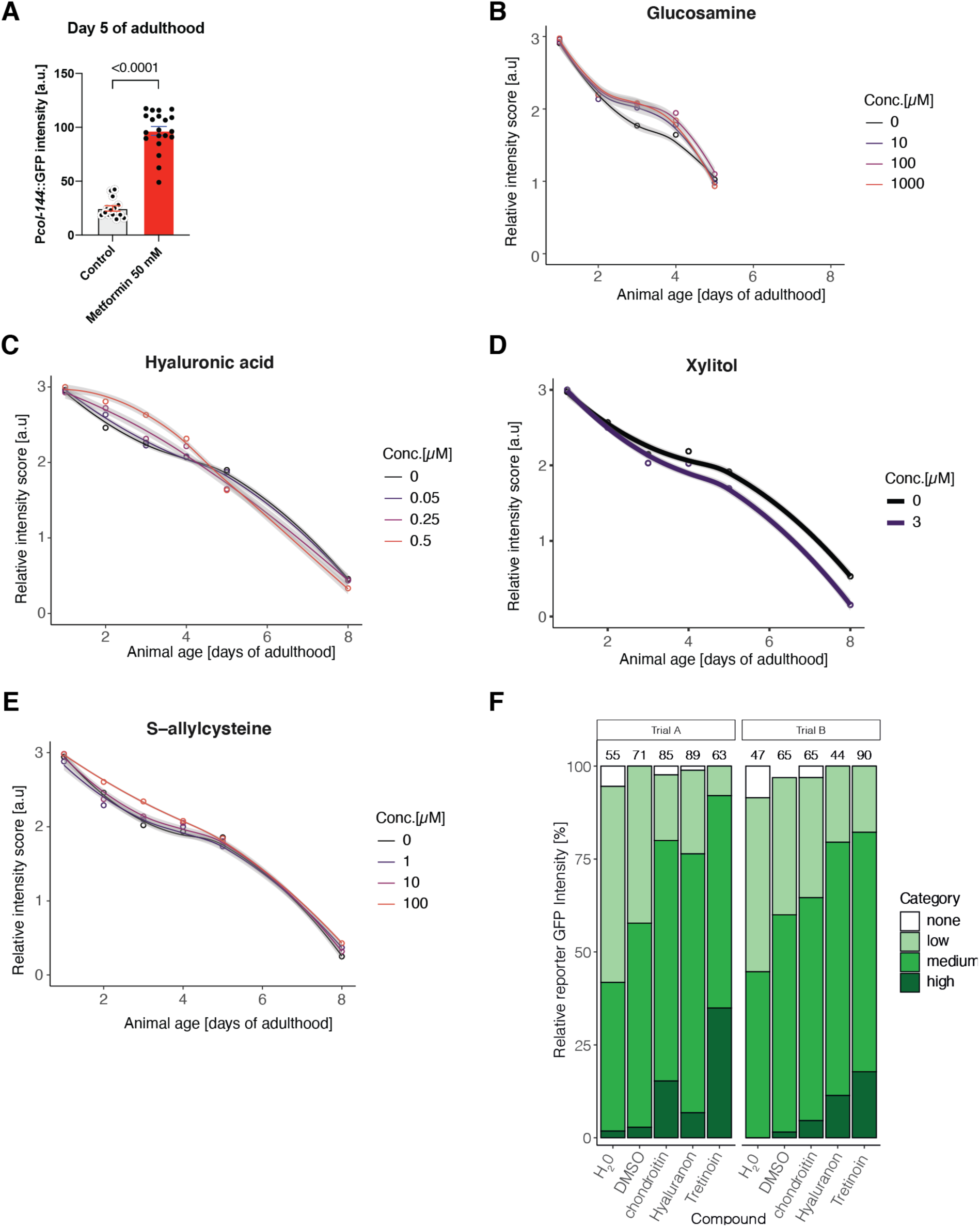
Screening compounds using prolonged collagen expression as a readout. (A) Quantification of collagen *col-144* promoter driven GFP (LSD2002 [P*col-144*::GFP]) upon metformin treatment started at day-2 of adulthood and scored at day-5 of adulthood. (B-E) Determining P*col-144*::GFP during aging for different compounds and concentrations. (F) Chondroitin, hyaluronic acid, or tretinoin treatment upon adulthood increased P*col-144*::GFP expression at day-5 of adulthood. For raw data, details, and statistics for (A-F), see Supplementary Table 6.

**Supplemental Figure 11:**
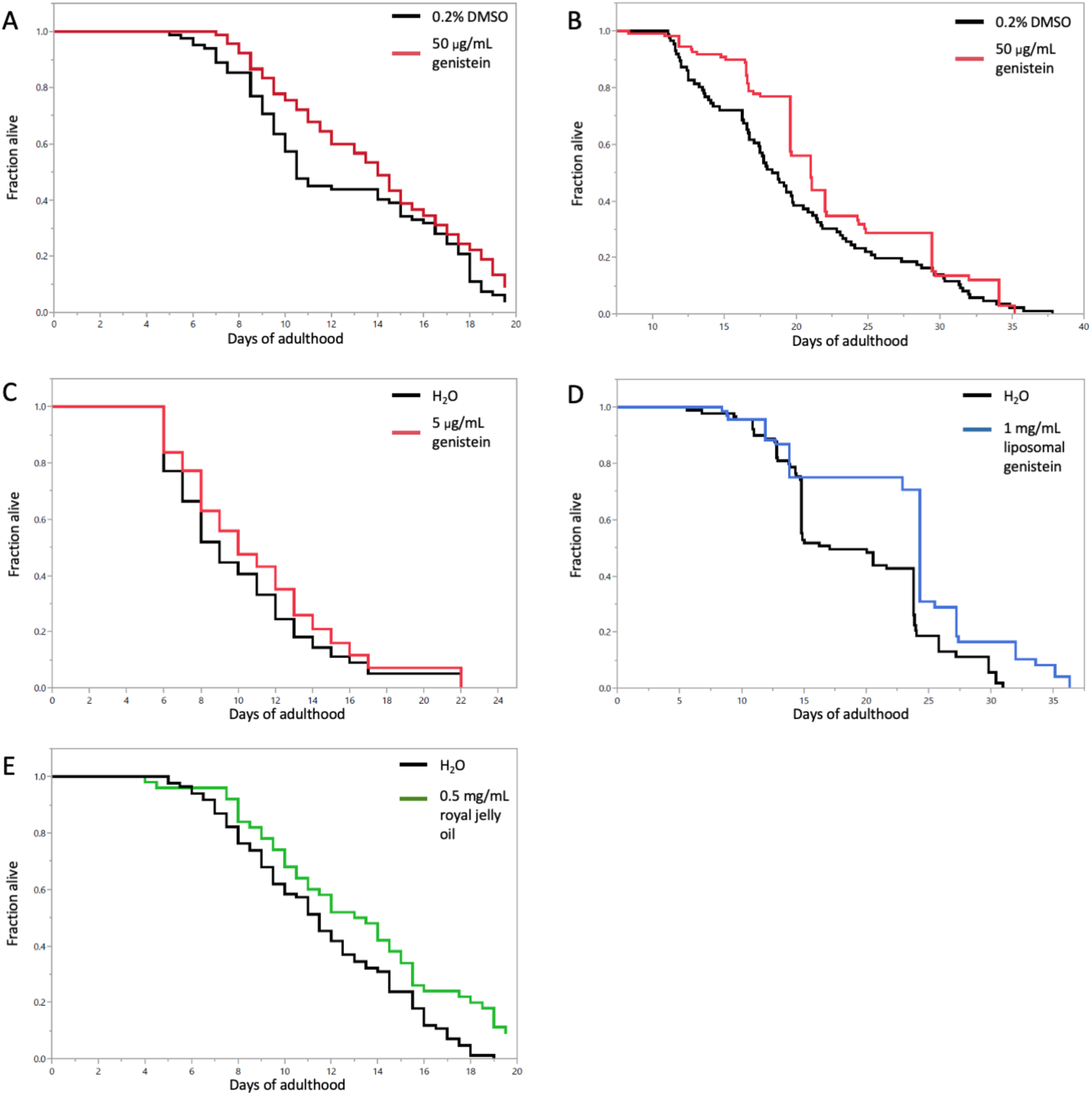
Lifespan extending compounds. Genistein extended lifespan in *C. elegans* at various concentrations, also when encapsulated in liposomes. (A) Genistein 50 μg/mL 11% mean lifespan extension compared to DMSO 0.2% [*P*-value 0.0497]. (B) Genistein 50 μg/mL 12.9% mean lifespan extension compared to DMSO 0.2% [*P* -value 0.0165]. (C) Genistein 5 μg/mL 9.8% mean lifespan extension compared to H_2_O [*P* -value 0.0297]. (D) Liposomal genistein 1 mg/mL 20.9% mean lifespan extension compared to H_2_O [*P*-value <.0001]. (E) Royal jelly oil 0.5 mg/mL 11% mean lifespan extension compared to H_2_O 0.2% [*P*-value 0.0497]. For raw data, detailed statistics and additional trials for (A-E), see Supplementary Table 7.

**Supplementary Figure 12.**
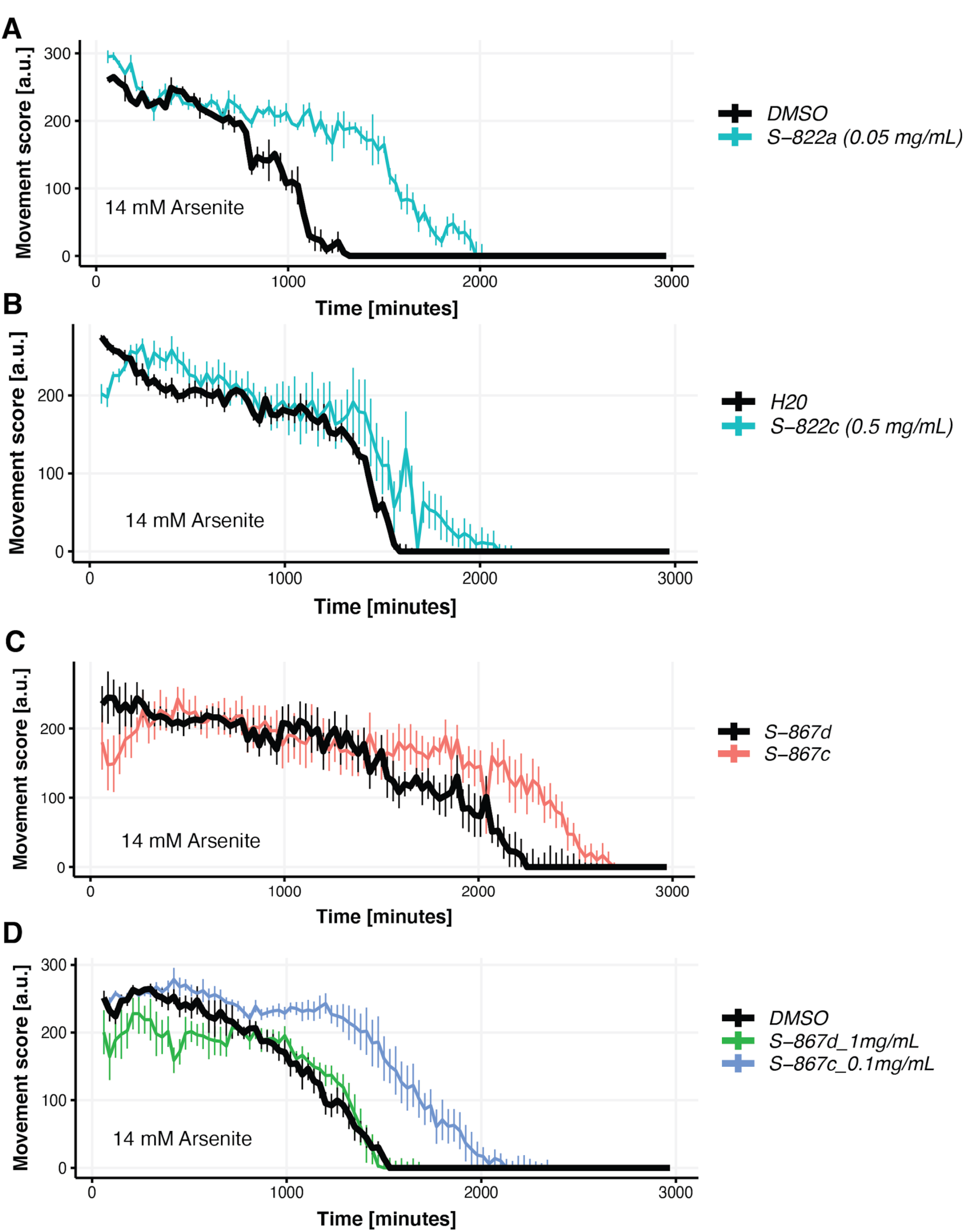
Genistein increases oxidative stress resistance. (A) Temperature-sensitive sterile wild-type background *C. elegans* (TJ1060) were fed S-822a genistein 0.05 mg/mL from day-1 of adulthood and at day-5 of adulthood tested for oxidative stress resistance. Genistein treated *C. elegans* survived longer in 14 mM arsenite. (B) Wild type (N2) animals were fed S-822c liposomal encapsulated genistein 0,5 mg/mL from L1 stage and tested for oxidative stress resistance at day-1 of adulthood. Genistein treated *C. elegans* survived longer in 14 mM arsenite. (C) Wild type (N2) animals were fed either S-867d liposomal vehicle or S-867c liposomal encapsulated genistein 0,5 mg/mL from L1 stage and tested for oxidative stress resistance at day-1 of adulthood. Genistein treated *C. elegans* survived longer in 14 mM arsenite. (D) Temperature-sensitive steril wild-type background *C. elegans* (TJ1060) were fed either S-867d liposomal vehicle or S-867c liposomal encapsulated genistein 0,5 mg/mL from day-1 of adulthood and at day-5 of adulthood tested for oxidative stress resistance. Genistein treated *C. elegans* survived longer in 14 mM arsenite. For raw data and statistics, see Supplementary Table 8.

## References

1. Aliper, A., Belikov, A.V., Garazha, A., Jellen, L., Artemov, A., Suntsova, M., Ivanova, A., Venkova, L., Borisov, N., Buzdin, A., et al. (2016). In search for geroprotectors: in silico screening and in vitro validation of signalome-level mimetics of young healthy state. Aging 8, 2127–2152.

2. Andreux, P.A., Blanco-Bose, W., Ryu, D., Burdet, F., Ibberson, M., Aebischer, P., Auwerx, J., Singh, A., and Rinsch, C. (2019). The mitophagy activator urolithin A is safe and induces a molecular signature of improved mitochondrial and cellular health in humans. Nature Metabolism 1, 595–603.

3. Bacalini, M.G., Boattini, A., Gentilini, D., Giampieri, E., Pirazzini, C., Giuliani, C., Fontanesi, E., Remondini, D., Capri, M., Rio, A.D., et al. (2015). A meta-analysis on age-associated changes in blood DNA methylation: results from an original analysis pipeline for Infinium 450k data. Aging 7, 97–109.

4. Baker-Williams, A.J., Hashmi, F., Budzyński, M.A., Woodford, M.R., Gleicher, S., Himanen, S.V., Makedon, A.M., Friedman, D., Cortes, S., Namek, S., et al. (2019). Co-chaperones TIMP2 and AHA1 Competitively Regulate Extracellular HSP90:Client MMP2 Activity and Matrix Proteolysis. Cell Reports 28, 1894–1906.e6.

5. Bakula, D., Aliper, A.M., Mamoshina, P., Petr, M.A., Teklu, A., Baur, J.A., Campisi, J., Ewald, C.Y., Georgievskaya, A., Gladyshev, V.N., et al. (2018). Aging and drug discovery. Aging 10, 3079–3088.

6. Bannister, C.A., Holden, S.E., Jenkins-Jones, S., Morgan, C.L., Halcox, J.P., Schernthaner, G., Mukherjee, J., and Currie, C.J. (2014). Can people with type 2 diabetes live longer than those without? A comparison of mortality in people initiated with metformin or sulphonylurea monotherapy and matched, non-diabetic controls. Diabetes, Obesity & Metabolism 16, 1165–1173.

7. Barardo, D., Thornton, D., Thoppil, H., Walsh, M., Sharifi, S., Ferreira, S., Anžič, A., Fernandes, M., Monteiro, P., Grum, T., et al. (2017a). The DrugAge database of aging-related drugs. Aging Cell 16, 594–597.

8. Barardo, D.G., Newby, D., Thornton, D., Ghafourian, T., Magalhães, J.P. de, and Freitas, A.A. (2017b). Machine learning for predicting lifespan-extending chemical compounds. Aging 9, 1721–1737.

9. Barzilai, N., Crandall, J.P., Kritchevsky, S.B., and Espeland, M.A. (2016). Metformin as a Tool to Target Aging. Cell Metabolism 23, 1060–1065.

10. Barzilai, N., Cuervo, A.M., and Austad, S. (2018). Aging as a Biological Target for Prevention and Therapy. JAMA: The Journal of the American Medical Association 320, 1321–1322.

11. Bell, G.A., Kantor, E.D., Lampe, J.W., Shen, D.D., and White, E. (2012a). Use of glucosamine and chondroitin in relation to mortality. European Journal of Epidemiology 27, 593–603.

12. Bell, J.T., Tsai, P.-C., Yang, T.-P., Pidsley, R., Nisbet, J., Glass, D., Mangino, M., Zhai, G., Zhang, F., Valdes, A., et al. (2012b). Epigenome-Wide Scans Identify Differentially Methylated Regions for Age and Age-Related Phenotypes in a Healthy Ageing Population. Plos Genet 8, e1002629.

13. Blankenburg, H., Pramstaller, P.P., and Domingues, F.S. (2018). A network-based meta-analysis for characterizing the genetic landscape of human aging. Biogerontology 19, 81–94.

14. Budovskaya, Y.V., Wu, K., Southworth, L.K., Jiang, M., Tedesco, P., Johnson, T.E., and Kim, S.K. (2008). An elt-3/elt-5/elt-6 GATA Transcription Circuit Guides Aging in C. elegans. Cell 134, 291–303.

15. Calvert, S., Tacutu, R., Sharifi, S., Teixeira, R., Ghosh, P., and Magalhães, J.P. de (2016). A network pharmacology approach reveals new candidate caloric restriction mimetics in C. elegans. Aging Cell 15, 256–266.

16. Campbell, J.M., Bellman, S.M., Stephenson, M.D., and Lisy, K. (2017). Metformin reduces all-cause mortality and diseases of ageing independent of its effect on diabetes control: A systematic review and meta-analysis. Ageing Research Reviews 40, 31–44.

17. Campisi, J., Kapahi, P., Lithgow, G.J., Melov, S., Newman, J.C., and Verdin, E. (2019). From discoveries in ageing research to therapeutics for healthy ageing. Nature 571, 183–192.

18. Chaturvedi, V., Kumar, J.U., Paithankar, K.R., Vanathi, P., and Sreedhar, A.S. (2011). Pharmacological Inhibition of Hsp90 as a Novel Antitumor Strategy to Target Cytoarchitecture Through Extracellular Matrix Signaling. Med Chem 7, 454–465.

19. Chen, G., Chen, H., Wang, C., Peng, Y., Sun, L., Liu, H., and Liu, F. (2012). Rapamycin ameliorates kidney fibrosis by inhibiting the activation of mTOR signaling in interstitial macrophages and myofibroblasts. Plos One 7, e33626.

20. Chung, C.L., Lawrence, I., Hoffman, M., Elgindi, D., Nadhan, K., Potnis, M., Jin, A., Sershon, C., Binnebose, R., Lorenzini, A., et al. (2019). Topical rapamycin reduces markers of senescence and aging in human skin: an exploratory, prospective, randomized trial. GeroScience 41, 861–869.

21. Consortium, Gte. (2013). The Genotype-Tissue Expression (GTEx) project. Nature Genetics 45, 580–585.

22. Dönertaş, H.M., Valenzuela, M.F., Partridge, L., and Thornton, J.M. (2018). Gene expression-based drug repurposing to target aging. Aging Cell 17, e12819.

23. Dönertaş, H.M., Fuentealba, M., Partridge, L., and Thornton, J.M. (2019). Identifying Potential Ageing-Modulating Drugs In Silico. Trends in Endocrinology and Metabolism: TEM 30, 118–131.

24. Espeland, M.A., Crimmins, E.M., Grossardt, B.R., Crandall, J.P., Gelfond, J.A.L., Harris, T.B., Kritchevsky, S.B., Manson, J.E., Robinson, J.G., Rocca, W.A., et al. (2017). Clinical Trials Targeting Aging and Age-Related Multimorbidity. The Journals of Gerontology Series A, Biological Sciences and Medical Sciences 72, 355–361.

25. Ewald, C.Y. (2019). The Matrisome during Aging and Longevity: A Systems-Level Approach toward Defining Matreotypes Promoting Healthy Aging. Gerontology 1–9.

26. Ewald, C.Y., Landis, J.N., Abate, J.P., Murphy, C.T., and Blackwell, T.K. (2015). Dauer-independent insulin/IGF-1-signalling implicates collagen remodelling in longevity. Nature 519, 97–101.

27. Ewald, C.Y., Marfil, V., and Li, C. (2016). Alzheimer-related protein APL-1 modulates lifespan through heterochronic gene regulation in Caenorhabditis elegans. Aging Cell 1–12.

28. Ewald, C.Y., Hourihan, J.M., and Blackwell, T.K. (2017). Oxidative Stress Assays (arsenite and tBHP) in Caenorhabditis elegans. BIO-PROTOCOL 7.

29. Fan, Y., Choi, T.-H., Chung, J.-H., Jeon, Y.-K., and Kim, S. (2019). Hyaluronic acid-cross-linked filler stimulates collagen type 1 and elastic fiber synthesis in skin through the TGF-β/Smad signaling pathway in a nude mouse model. J Plastic Reconstr Aesthetic Surg Jpras 72, 1355–1362.

30. Fernandes, M., Wan, C., Tacutu, R., Barardo, D., Rajput, A., Wang, J., Thoppil, H., Thornton, D., Yang, C., Freitas, A., et al. (2016). Systematic analysis of the gerontome reveals links between aging and age-related diseases. Human Molecular Genetics 25, 4804–4818.

31. Fisher, G.J., Quan, T., Purohit, T., Shao, Y., Cho, M.K., He, T., Varani, J., Kang, S., and Voorhees, J.J. (2009). Collagen fragmentation promotes oxidative stress and elevates matrix metalloproteinase-1 in fibroblasts in aged human skin. American Journal Of Pathology 174, 101–114.

32. Fuentealba, M., Dönertaş, H.M., Williams, R., Labbadia, J., Thornton, J.M., and Partridge, L. (2019). Using the drug-protein interactome to identify anti-ageing compounds for humans. PLoS Computational Biology 15, e1006639.

33. Griffiths, C., Russman, A.N., Majmudar, G., Singer, R.S., Hamilton, T.A., and Voorhees, J.J. (1993). Restoration of Collagen Formation in Photodamaged Human Skin by Tretinoin (Retinoic Acid). New Engl J Med 329, 530–535.

34. Gueniche, A., and Castiel-Higounenc, I. (2017). Efficacy of Glucosamine Sulphate in Skin Ageing: Results from an ex vivo Anti-Ageing Model and a Clinical Trial. Skin Pharmacol Physi 30, 36–41.

35. Haes, W.D., Frooninckx, L., Assche, R.V., Smolders, A., Depuydt, G., Billen, J., Braeckman, B.P., Schoofs, L., and Temmerman, L. (2014). Metformin promotes lifespan through mitohormesis via the peroxiredoxin PRDX-2. Proc National Acad Sci 111, E2501–E2509.

36. Hunter, M.C., O’Hagan, K.L., Kenyon, A., Dhanani, K.C.H., Prinsloo, E., and Edkins, A.L. (2014). Hsp90 binds directly to fibronectin (FN) and inhibition reduces the extracellular fibronectin matrix in breast cancer cells. Plos One 9, e86842.

37. Ihaka, R., and Gentleman, R. (2012). R: A Language for Data Analysis and Graphics. J Comput Graph Stat 5, 299–314.

38. Janssens, G.E., Lin, X.-X., Millan-Ariño, L., Kavšek, A., Sen, I., Seinstra, R.I., Stroustrup, N., Nollen, E.A.A., and Riedel, C.G. (2019). Transcriptomics-Based Screening Identifies Pharmacological Inhibition of Hsp90 as a Means to Defer Aging. Cell Reports 27, 467–480.e6.

39. Kennedy, B.K., Berger, S.L., Brunet, A., Campisi, J., Cuervo, A.M., Epel, E.S., Franceschi, C., Lithgow, G.J., Morimoto, R.I., Pessin, J.E., et al. (2014). Geroscience: linking aging to chronic disease. Cell 159, 709–713.

40. Kim, E., Aging, S.L.T.M. of, and 2019 (2019). Recent progresses on anti-aging compounds and their targets in Caenorhabditis elegans. Elsevier 3, 121–124.

41. Komljenovic, A., Li, H., Sorrentino, V., Kutalik, Z., Auwerx, J., and Robinson-Rechavi, M. (2019). Cross-species functional modules link proteostasis to human normal aging. Plos Comput Biol 15, e1007162.

42. Lamb, J., Crawford, E.D., Peck, D., Modell, J.W., Blat, I.C., Wrobel, M.J., Lerner, J., Brunet, J.-P., Subramanian, A., Ross, K.N., et al. (2006). The Connectivity Map: using gene-expression signatures to connect small molecules, genes, and disease. Science (New York, NY) 313, 1929–1935.

43. Lampi, M.C., and Reinhart-King, C.A. (2018). Targeting extracellular matrix stiffness to attenuate disease: From molecular mechanisms to clinical trials. Science Translational Medicine 10, eaao0475.

44. Lee, E.B., Ahn, D., Kim, B.J., Lee, S.Y., Seo, H.W., Cha, Y.-S., Jeon, H., Eun, J.S., Cha, D.S., and Kim, D.K. (2015). Genistein from Vigna angularis Extends Lifespan in Caenorhabditis elegans. Biomol Ther 23, 77–83.

45. Lephart, E.D., and Andrus, M.B. (2017). Human skin gene expression: Natural (trans) resveratrol versus five resveratrol analogs for dermal applications. Experimental Biology and Medicine (Maywood, N.J.) 242, 1482–1489.

46. Li, P., Liang, M.-L., Zhu, Y., Gong, Y.-Y., Wang, Y., Heng, D., and Lin, L. (2014). Resveratrol inhibits collagen I synthesis by suppressing IGF-1R activation in intestinal fibroblasts. World J Gastroentero 20, 4648.

47. Li, Z.-H., Gao, X., Chung, V.C., Zhong, W.-F., Fu, Q., Lv, Y.-B., Wang, Z.-H., Shen, D., Zhang, X.-R., Zhang, P.-D., et al. (2020). Associations of regular glucosamine use with all-cause and cause-specific mortality: a large prospective cohort study. Ann Rheum Dis 79, annrheumdis-2020-217176.

48. Lippiello, L. (2007). Collagen Synthesis in Tenocytes, Ligament Cells and Chondrocytes Exposed to a Combination of Glucosamine HCl and Chondroitin Sulfate. Evid-Based Compl Alt 4, 219–224.

49. Liu, H., Guo, M., Xue, T., Guan, J., Luo, L., and Zhuang, Z. (2016). Screening lifespan-extending drugs in Caenorhabditis elegans via label propagation on drug-protein networks. BMC Systems Biology 10, 131.

50. López-Otín, C., Blasco, M.A., Partridge, L., Serrano, M., and Kroemer, G. (2013). The hallmarks of aging. Cell 153, 1194–1217.

51. Lu, Y., Brommer, B., Tian, X., Krishnan, A., Meer, M., Wang, C., Vera, D.L., Zeng, Q., Yu, D., Bonkowski, M.S., et al. (2020). Reprogramming to recover youthful epigenetic information and restore vision. Nature 588, 124–129.

52. Lucanic, M., Lithgow, G.J., and Alavez, S. (2013). Pharmacological lifespan extension of invertebrates. Ageing Res Rev 12, 445–458.

53. Magalhães, J.P. de, and Toussaint, O. (2004). GenAge: a genomic and proteomic network map of human ageing. Febs Lett 571, 243–247.

54. Magalhães, J.P. de, Curado, J., and Church, G.M. (2009). Meta-analysis of age-related gene expression profiles identifies common signatures of aging. Bioinform Oxf Engl 25, 875–881.

55. Mannick, J.B., Giudice, G.D., Lattanzi, M., Valiante, N.M., Praestgaard, J., Huang, B., Lonetto, M.A., Maecker, H.T., Kovarik, J., Carson, S., et al. (2014). mTOR inhibition improves immune function in the elderly. Science Translational Medicine 6, 268ra179–268ra179.

56. Mannick, J.B., Morris, M., Hockey, H.-U.P., Roma, G., Beibel, M., Kulmatycki, K., Watkins, M., Shavlakadze, T., Zhou, W., Quinn, D., et al. (2018). TORC1 inhibition enhances immune function and reduces infections in the elderly. Science Translational Medicine 10, eaaq1564.

57. Matori, H., Umar, S., Nadadur, R.D., Sharma, S., Partow-Navid, R., Afkhami, M., Amjedi, M., and Eghbali, M. (2012). Genistein, a Soy Phytoestrogen, Reverses Severe Pulmonary Hypertension and Prevents Right Heart Failure in Rats. Hypertension 60, 425–430.

58. Moskalev, A., Chernyagina, E., Magalhães, J.P. de, Barardo, D., Thoppil, H., Shaposhnikov, M., Budovsky, A., Fraifeld, V.E., Garazha, A., Tsvetkov, V., et al. (2015). Geroprotectors.org: a new, structured and curated database of current therapeutic interventions in aging and age-related disease. Aging 7, 616–628.

59. Naba, A., Clauser, K.R., Ding, H., Whittaker, C.A., Carr, S.A., and Hynes, R.O. (2016). The extracellular matrix: Tools and insights for the “omics” era. Matrix Biology: Journal of the International Society for Matrix Biology 49, 10–24.

60. Niccoli, T., and Partridge, L. (2012). Ageing as a risk factor for disease. Current Biology: CB 22, R741–52.

61. Nkuipou-Kenfack, E., Bhat, A., Klein, J., Jankowski, V., Mullen, W., Vlahou, A., Dakna, M., Koeck, T., Schanstra, J.P., Zürbig, P., et al. (2015). Identification of ageing-associated naturally occurring peptides in human urine. Oncotarget 6, 34106–34117.

62. Olshansky, S.J. (2018). From Lifespan to Healthspan. JAMA: The Journal of the American Medical Association 320, 1323–1324.

63. Partridge, L., Deelen, J., and Slagboom, P.E. (2018). Facing up to the global challenges of ageing. Nature 561, 45–56.

64. Petrascheck, M., Ye, X., and Buck, L.B. (2007). An antidepressant that extends lifespan in adult Caenorhabditis elegans. Nature 450, 553–556.

65. Polito, F., Marini, H., Bitto, A., Irrera, N., Vaccaro, M., Adamo, E.B., Micali, A., Squadrito, F., Minutoli, L., and Altavilla, D. (2012). Genistein aglycone, a soy-derived isoflavone, improves skin changes induced by ovariectomy in rats: Genistein improves skin changes in OVX rats. Brit J Pharmacol 165, 994–1005.

66. Pryor, R., and Cabreiro, F. (2015). Repurposing metformin: an old drug with new tricks in its binding pockets. Biochem J 471, 307–322.

67. Riera, C.E., and Dillin, A. (2015). Can aging be “drugged”? Nature Medicine 21, 1400–1405.

68. Ristow, M., and Schmeisser, K. (2014). Mitohormesis: Promoting Health and Lifespan by Increased Levels of Reactive Oxygen Species (ROS). Dose-Response: A Publication of International Hormesis Society 12, 288–341.

69. Sathyan, S., Ayers, E., Gao, T., Weiss, E.F., Milman, S., Verghese, J., and Barzilai, N. (2020). Plasma proteomic profile of age, health span, and all-cause mortality in older adults. Aging Cell 19, e13250.

70. Socovich, A.M., and Naba, A. (2019). The cancer matrisome: From comprehensive characterization to biomarker discovery. Seminars in Cell & Developmental Biology 89, 157–166.

71. Sood, S., Gallagher, I.J., Lunnon, K., Rullman, E., Keohane, A., Crossland, H., Phillips, B.E., Cederholm, T., Jensen, T., Loon, L.J.C. van, et al. (2015). A novel multi-tissue RNA diagnostic of healthy ageing relates to cognitive health status. Genome Biol 16, 185.

72. Statzer, C., and Ewald, C.Y. (2020). The extracellular matrix phenome across species. Matrix Biology Plus 8, 100039.

73. Stroustrup, N., Ulmschneider, B.E., Nash, Z.M., López-Moyado, I.F., Apfeld, J., and Fontana, W. (2013). The Caenorhabditis elegans Lifespan Machine. Nature Methods 10, 665–670.

74. Stroustrup, N., Anthony, W.E., Nash, Z.M., Gowda, V., Gomez, A., López-Moyado, I.F., Apfeld, J., and Fontana, W. (2016). The temporal scaling of Caenorhabditis elegans ageing. Nature 530, 103–107.

75. Taha, I.N., and Naba, A. (2019). Exploring the extracellular matrix in health and disease using proteomics. Essays in Biochemistry 63, 417–432.

76. Tarkhov, A.E., Alla, R., Ayyadevara, S., Pyatnitskiy, M., Menshikov, L.I., Reis, R.J.S., and Fedichev, P.O. (2019). A universal transcriptomic signature of age reveals the temporal scaling of Caenorhabditis elegans aging trajectories. Scientific Reports 9, 7368.

77. Teuscher, A.C., and Ewald, C.Y. (2018). Overcoming Autofluorescence to Assess GFP Expression During Normal Physiology and Aging in Caenorhabditis elegans. BIO-PROTOCOL 8.

78. Teuscher, A.C., Jongsma, E., Davis, M.N., Statzer, C., Gebauer, J.M., Naba, A., and Ewald, C.Y. (2019a). The in-silico characterization of the Caenorhabditis elegans matrisome and proposal of a novel collagen classification. Matrix Biology Plus 1–13.

79. Teuscher, A.C., Statzer, C., Pantasis, S., Bordoli, M.R., and Ewald, C.Y. (2019b). Assessing Collagen Deposition During Aging in Mammalian Tissue and in Caenorhabditis elegans. Methods in Molecular Biology (Clifton, NJ) 1944, 169–188.

80. Therneau, T.M., and Grambsch, P.M. (2000). Modeling Survival Data: Extending the Cox Model. 1–6.

81. Tsubota, K. (2016). The first human clinical study for NMN has started in Japan. Npj Aging and Mechanisms of Disease 2, 16021.

82. Tyshkovskiy, A., Bozaykut, P., Borodinova, A.A., Gerashchenko, M.V., Ables, G.P., Garratt, M., Khaitovich, P., Clish, C.B., Miller, R.A., and Gladyshev, V.N. (2019). Identification and Application of Gene Expression Signatures Associated with Lifespan Extension. Cell Metabolism 30, 573–593.e8.

83. Vanhaelen, Q., Mamoshina, P., Aliper, A.M., Artemov, A., Lezhnina, K., Ozerov, I., Labat, I., and Zhavoronkov, A. (2017). Design of efficient computational workflows for in silico drug repurposing. Drug Discov Today 22, 210–222.

84. Wang, F., Garza, L.A., Kang, S., Varani, J., Orringer, J.S., Fisher, G.J., and Voorhees, J.J. (2007). In Vivo Stimulation of De Novo Collagen Production Caused by Cross-linked Hyaluronic Acid Dermal Filler Injections in Photodamaged Human Skin. Arch Dermatol 143, 155–163.

85. Wang, J., Zhang, S., Wang, Y., Chen, L., and Zhang, X.-S. (2009a). Disease-aging network reveals significant roles of aging genes in connecting genetic diseases. Plos Comput Biol 5, e1000521.

86. Wang, X., Zhao, Y., Wong, K., Ehlers, P., Kohara, Y., Jones, S.J., Marra, M.A., Holt, R.A., Moerman, D.G., and Hansen, D. (2009b). Identification of genes expressed in the hermaphrodite germ line of C. elegans using SAGE. BMC Genomics 10, 213.

87. Weimer, S., Priebs, J., Kuhlow, D., Groth, M., Priebe, S., Mansfeld, J., Merry, T.L., Dubuis, S., Laube, B., Pfeiffer, A.F., et al. (2014). D-Glucosamine supplementation extends life span of nematodes and of ageing mice. Nature Communications 5, 3563.

88. Wick, G., Grundtman, C., Mayerl, C., Wimpissinger, T.-F., Feichtinger, J., Zelger, B., Sgonc, R., and Wolfram, D. (2013). The Immunology of Fibrosis. Annual Review of Immunology, Vol 31 31, 107–135.

89. Wynn, T. (2008). Cellular and molecular mechanisms of fibrosis. J Pathology 214, 199–210.

90. Yang, J., Huang, T., Petralia, F., Long, Q., Zhang, B., Argmann, C., Zhao, Y., Mobbs, C.V., Schadt, E.E., Zhu, J., et al. (2015). Synchronized age-related gene expression changes across multiple tissues in human and the link to complex diseases. Sci Rep-Uk 5, 15145.

91. Yang, J., Peng, S., Zhang, B., Houten, S., Schadt, E., Zhu, J., Suh, Y., and Tu, Z. (2020). Human geroprotector discovery by targeting the converging subnetworks of aging and age-related diseases. Geroscience 42, 353–372.

92. Ye, X., Linton, J.M., Schork, N.J., Buck, L.B., and Petrascheck, M. (2014). A pharmacological network for lifespan extension in Caenorhabditis elegans. Aging Cell 13, 206–215.

93. Zeng, L., Yang, J., Peng, S., Zhu, J., Zhang, B., Suh, Y., and Tu, Z. (2020). Transcriptome analysis reveals the difference between “healthy” and “common” aging and their connection with age-related diseases. Aging Cell 19, e13121.

94. Zhavoronkov, A., Buzdin, A.A., Garazha, A.V., Borisov, N.M., and Moskalev, A.A. (2014). Signaling pathway cloud regulation for in silico screening and ranking of the potential geroprotective drugs. Frontiers in Genetics 5, 49.

